# comBO: A combined human bone and lympho-myeloid bone marrow organoid for pre-clinical modelling of haematopoietic disorders

**DOI:** 10.1101/2025.02.16.638505

**Authors:** Yuqi Shen, Camelia Benlabiod, Antonio Rodriguez-Romera, Edmund Watson, Jasmeet S. Reyat, Kristian Gurashi, Shady Adnan-Awad, Rupen Hargreaves, Samuel Kemble, Charlotte G. Smith, Adam P. Croft, Udo Opperman, Natalie Jooss, Zoe C. Wong, Julie Rayes, Adam J. Mead, Anindita Roy, Sarah Gooding, Bethan Psaila, Abdullah O. Khan

## Abstract

The bone marrow supports lifelong blood and immune cell production. Current human bone marrow organoid models do not include both lymphoid and myeloid elements and lack the complexity of stromal cell types present in native haematopoietic tissues, precluding the accurate *ex vivo* modelling of human pathologies. Here we introduce “comBOs” (**<u>com</u>**bined **b**one and lympho-myeloid bone marrow **o**rganoids) that include osteolineage, vascular, lymphoid and myeloid cells. comBOs are generated by the differentiation of induced pluripotent stem cells guided by physiologically-relevant oxygen and cytokine exposures within an innovative granular microgel scaffold to increase scalability and reproducibility. We demonstrate that comBOs can be used to generate “chimeroids” - incorporating healthy or aberrant cells from adult donors – and recapitulate features of diseased microenvironments. ComBOs are one of the most physiologically-relevant human organoid systems to date, and this study showcases the potential of 3D *in vitro* disease models for discovery science and translational studies.

## Introduction

The bone marrow is the primary site of post-natal haematopoiesis and is organised into specialized niches that support blood and immune cell production^1,2^. An elaborate vasculature and diverse stromal cell types regulate haematopoietic stem and progenitor cells (HSPCs), guiding differentiation, quiescence, and expansion^3–5^. The stromal repertoire of the bone marrow includes mesenchymal stem cells (MSCs), which generate osteo- and adipo-lineage cells, as well as specific MSC subsets (e.g. Leptin Receptor (LEPR+) and Nestin (NES+) MSCs)^1,4,6^. These extrinsic cues maintain blood cell production at steady state, and in response to physiological challenges.

Disruption of the bone marrow microenvironment contributes to the pathogenesis of bone marrow disorders, including blood malignancies and bone marrow failure syndromes^7^. Disease-mediated changes in the microenvironment, such as inflammation, are known to limit the efficacy of therapies, and contribute to chemoresistance and relapse^7,8^. Similarly, age-associated changes to the niche contribute to the myeloid-biased haematopoiesis and immune decline that occurs during human ageing^9^.

Human bone marrow is a relatively inaccessible organ, and important species-specific differences in haematopoiesis between humans and mice the translatability of animal disease models^10–12^. Therefore, human *in vitro* tissue systems that faithfully capture bone marrow niches are required for the accurate mechanistic study of human pathologies. Recent advances in the development of human bone marrow organoids have begun to address this gap^13–15^. We, and others, have recently developed myeloid bone marrow organoids (mBMOs) comprised of myelopoietic cells, endothelium, fibroblasts and MSCs^16,17^, and demonstrated their utility in modelling myeloid disorders and genetic haematopathologies^18,19^. Despite these advances, current models have three key limitations: they do not capture simultaneous myelopoiesis and lymphopoiesis, they lack osteoblast-lineage cells and bone deposition potential, and the scalability and reproducibility is limited by the requirement for embedding and extraction from hydrogel scaffolds.

Here, we describe the generation of **com**bined **b**one and lympho-myeloid **b**one marrow **o**rganoids (comBOs), a highly scalable, physiologically relevant multi-niche model of the human bone marrow.

ComBOs generate both lymphoid and myeloid lineage cells and contain a complexity of stromal cell types including osteolineage, adipocyte and MSC subtypes which demonstrate transcriptional homology to adult bone marrow. We developed the use of granular microgel scaffolds, which greatly improve the scalability and reproducibility of organoids compared to previous bulk hydrogel approaches, enabling higher-throughput organoid production for discovery research and preclinical applications.

We demonstrate the utility of comBOs to model human disease by engrafting cells from patients with multiple myeloma, a plasma cell malignancy in which inflammatory remodelling of the bone marrow niche and bony destruction are key hallmarks of disease^20,21^. Human myeloma comBO ‘chimeroids’ faithfully capture myeloma-induced stromal inflammation and metabolic reprogramming of MSCs as well as arrested maturation of osteoblasts. Dissecting receptor-ligand interactions between myeloma cells and organoid niche components identified macrophage inhibitory factor (MIF) as a central mediator of stromal inflammation, and we show that inhibition of MIF mitigates the inflammatory responses induced by primary myeloma cells.

ComBOs represent an advanced, scalable, multi-niche human organoid model that advances the field of *in vitro* human tissue modelling by providing a robust platform to interrogate bone marrow pathologies. More broadly, comBO is one of the most faithful human tissue systems reported to date, showcasing the potential of *in vitro* iPSC (induced pluripotent stem cell)-derived 3D systems to emulate native tissues.

## Results

### Development of comBO: a single differentiation generating lympho-myeloid, vascular, and osteogenic cell types

To generate a 3D culture capturing lympho-myeloid and bone marrow stromal elements, human iPSCs were subjected to a series of cytokine cocktails and hydrogel embedding to guide multi-lineage differentiation at physiological oxygen level (5% O_2_) **(Fig 1a, Supp Fig 1**)^22^. This involved mesodermal induction prior to commitment to vascular and myeloid lineages, followed by a lymphoid phase where myeloid factors were depleted and the lymphopoietic cytokine IL7 was supplemented. Finally, individual organoids were extracted from a hydrogel scaffold for downstream culture (**Supp Fig 1**)^23^.

**Figure 1:**
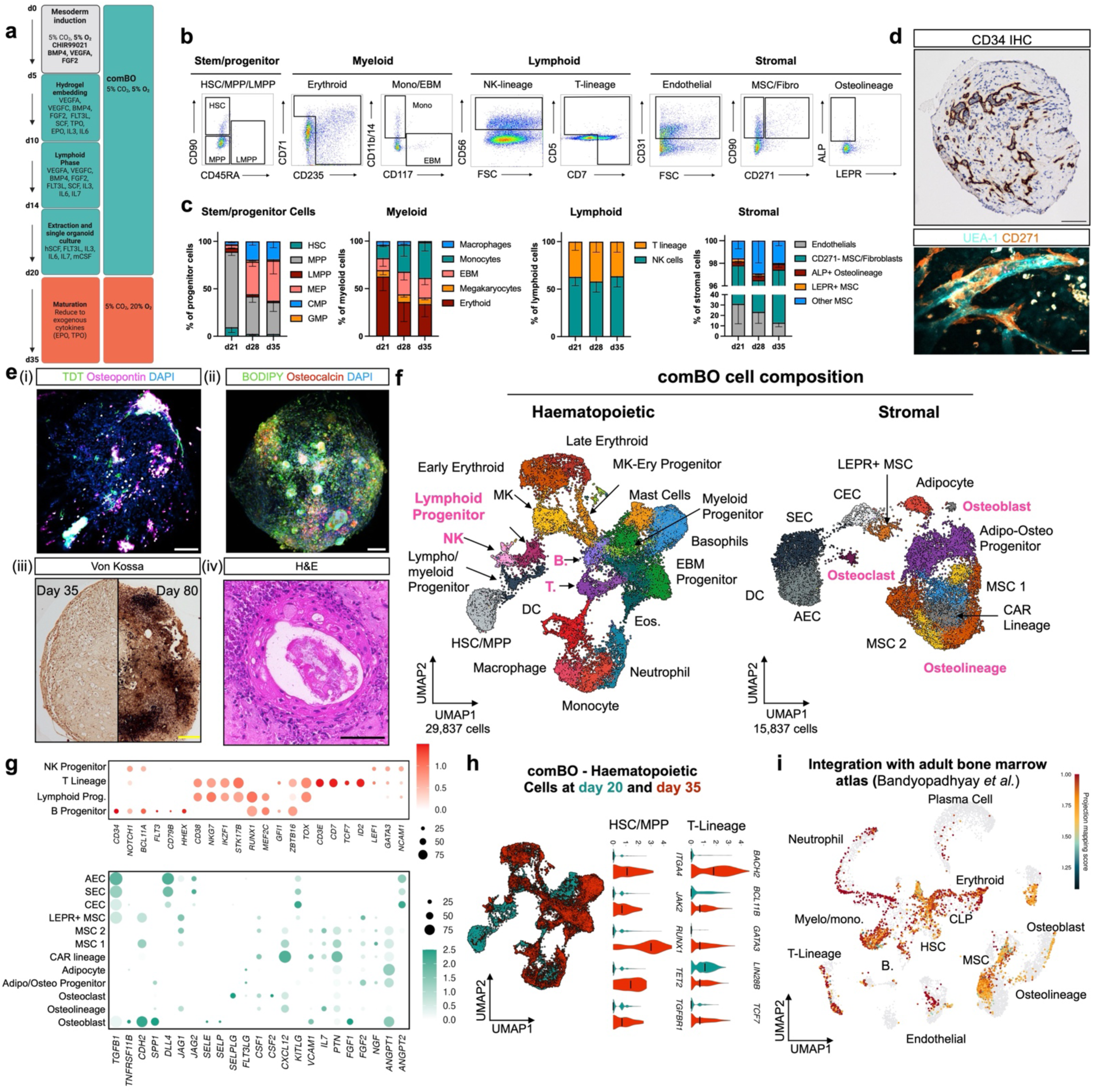
Development and validation of a protocol to generate an iPSC-derived combined bone and lympho-myeloid bone marrow organoid (comBO). **(a)** Schematic of workflow. ComBOs are generated in bone marrow relevant oxygen concentration (5% O2). **(b)** Representative flow plots of key comBO haematopoietic and stromal populations at day 21. **(c)** Quantified relative percentages of haematopoietic and stromal cell types in each flow panel over time (*n =* 4 separate differentiations). Each cell type was identified and quantified using the gating strategy shown in Supp Fig 2 and 3, followed by the calculation of its relative proportion within the defined populations. (Error bars are standard deviation). **(d)** Validation of vascular architecture using CD34 staining (scale bar = 100 μm) and confocal microscopy (vasculature (UEA-1), perivascular MSC (CD271+), scale bar = 25 μm). **(e)** Imaging to confirm early (i) lymphoid (TDT+) cells and early osteogenesis (osteopontin+) (scale bar = 100 μm) (ii) Adipocytes identified using BODIPY+ (scale bar = 100 μm). (iii) Von Kossa staining (scale bar = 200 μm) and (iv) H&E staining to confirm bone formation by day 80 (scale bar = 50 μm). **(f)** UMAP of single cell data characterising the haematopoietic and stromal cells generated at day 20 and day 35 by comBOs to transcriptionally profile cells and assess maturation **(g)** Bubble plots characterising gene expression in lymphoid populations (top), and the expression of key bone marrow haematopoietic factors by stromal cells (bottom). **(h)** Comparison of day 20 and day 35 comBO transcriptome highlighting canonical changes in the expression of adult genes in HSC/MPP and T-lineage. (i) Integration with a recently published atlas of adult human bone marrow revealing homology and overlap with cells of the adult bone marrow. (*n* = 4 separate differentiations, 24 organoids pooled per repeat. Schematic generated on Biorender.com).

This protocol generated HSPCs, myeloid cells (erythroid, megakaryocyte, monocyte, and eosinophil/basophil/mast (EBM) cells), lymphoid cells (T-lineage and natural killer (NK) cells), mesenchymal stromal, osteolineage and endothelial cells at day 21 of culture **(Fig 1b, Supp Fig 2, Supp Table 1)**. A further two-week culture at 20% O_2_ allowed for the maturation of key populations. T-lineage cells, for example, are CD7+ CD5- at day 21, and become CD5+ CD7+ on day 35 of culture **(Supp Fig 3a-b)**.

Imaging confirmed a lumen-forming, CD34+ vascular network (**Fig 1d**), with vessels supported by CD271+ perivascular cells (**Fig 1d**). Osteopontin and TDT (Terminal deoxynucleotidyl transferase) positive cells were detected, confirming the presence of osteo- and T-lymphoid lineage cells (**Fig 1e(i)**), as well as BODIPY+ adipocytes (**Fig 1e(ii)**). Remarkably, progressive mineralization was observed over time (Von Kossa, **Fig 1e(iii). Supp Fig 3c**), confirming the presence of osteolineage cells with bone-depositing capacity (**Fig 1e(iv)**). We confirmed the clonogenic potential of HSPCs vis Colony Forming Unit (CFU) assay using comBO-derived cells compared to peripheral blood controls (**Supp Fig 7**). Together, these data confirm generation of lymphoid, myeloid, vascular and mesenchymal stromal including functional osteogenesis and adipogenesis within a single human organoid system.

### ComBOs show transcriptional homology to adult human bone marrow

To investigate the homology of the organoid cell types to those present in human bone marrow, single cell RNA sequencing (scRNA-seq) was performed on organoids at two timepoints (days 20 and 35). Together, cells from 4 independent differentiations were pooled to generate a data set of 29,333 haematopoietic and 15,837 stromal cells (**Fig 1f**). Cell types were annotated based on canonical gene expression (**Fig 1f, Supp Fig 4,5**)^2,24,25^.

The haematopoietic compartment included mono/macrophage cells, neutrophils dendritic cells, lymphoid and myeloid progenitors, T-progenitors, eo/baso/mast cells, HSPCs, erythroid cells and megakaryocytes (**Fig 1f, Supp Fig 3**). The stromal compartment was comprised of endothelial cells (sinusoidal (SEC), arterial (AEC), and capillary (CEC) endothelial cells), MSCs, CAR (CXCL12 abundant reticular cell) progenitors, adipo/osteo lineage cells (including osteoblasts), and fibroblasts (**Fig 1f, Supp Fig 5**).

comBO demonstrated, for the first time, the generation of lymphoid populations in parallel to myeloid cells in a single organoid differentiation^16,18,26^. These included B-progenitor (*CD34, CD79B, FLT3, BCL11A*), lymphoid progenitor (*CD38, NKG7, IKZF1, RUNX1, MEF2C, TOX)*, T-lineage (*CD3E, CD7, TCF7),* and NK progenitor (*LEF1, GATA3, NCAM1*) populations (**Fig 1g, Supp Fig 5**). Similarly, osteoblasts (*POSTN, SPP1, ANGPT1, KITLG, KLL1, WNT5A, SP7, BGLAP*), osteolineage progenitors *(BMP2, ALPL, JAG1, GREM1, CDH11),* and osteoclasts *(MMP9, TNFRS11A. TNFSF11)* were identified (**Supp Fig 5**). Bone marrow stromal cells showed expression of key growth factors, chemokines and adhesion factors including *KITLG, TGFB1, DLL4, JAG1/2, CXCL12, IL7, ANGPT1/2*, confirming the production of haematopoietic support factors by the organoid niche components (**Fig 1g**).

A comparison of transcriptomic data acquired at day 20 and day 35 demonstrated the differentiation of the cell types observed, with an increase in the expression of markers of adult haematopoiesis indicating a maturation of comBO haematopoietic cells (**Fig 1h, Supp Fig 6a-b**). HSC/MPPs showed a notable increase in *ITGA4, JAK2, RUNX1, TET2* and *TGFBR1*, and T-lineage cells demonstrated an increase in *BACH2, BCL11B, GATA3* and *TCF7* and a decrease in fetal *LIN28B* over time (**Fig 1h**).

We integrated our day 35 data with a recently published adult human bone marrow atlas and found a high degree of transcriptional homology between comBO cells and adult bone marrow cells (**Fig 1i, Supp Fig 8**)^2^.

### Using granular microgels enables high-throughput organoid generation

The use of ‘bulk’ hydrogels creates limitations to scalability and cost due to labour-intensive preparation, the requirement for manual extraction and replating of organoids which preclude automation for higher-throughput pharmacogenomic studies^16–18^.

Inspired by the use of granular microgels—assemblies of discrete microgel particles that can be physically compacted into a support structure—for 3D bioprinting, we developed a workflow to overcome the current limitations of bulk hydrogel embedding^27–30^. Hydrogels were mechanically processed to generate a granular microgel (**Fig 2a, Supp Fig 9**), then combined with media, growth factors and cell aggregates and dispensed into wells of 96- (or 384-) well ultra-low attachment plates followed by centrifugation to compact the microgel (**Fig 2a**)^28^. We found that the compacted microgels supported vascular sprouting in a similar manner to bulk hydrogels whilst using a fraction of the material (80% reduction in bulk gel volume) (**Fig 2b**). Organoids generated in this way do not require manual extraction, as excess microgel is washed away in subsequent media changes.

**Figure 2:**
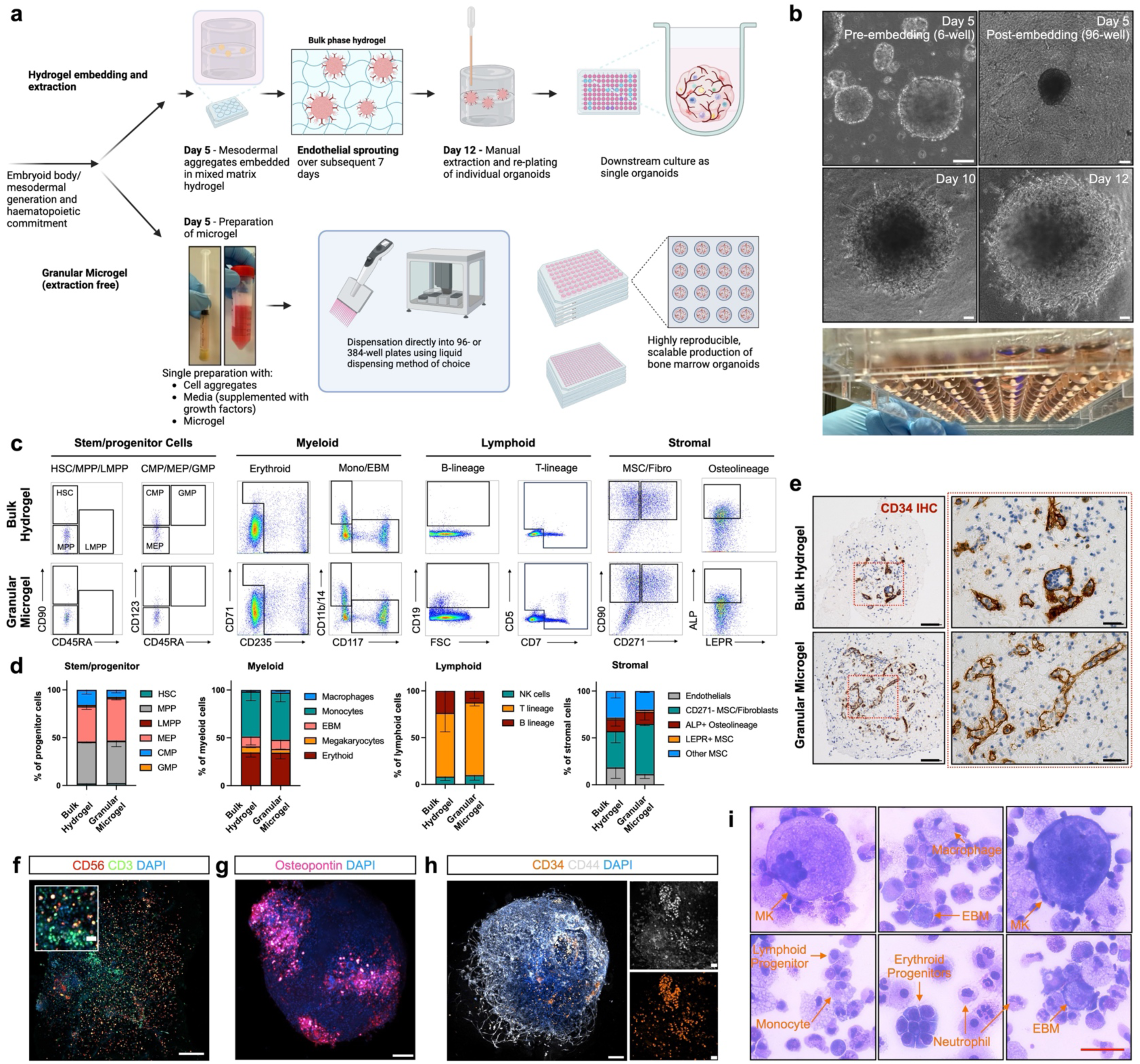
Development and validation of a high throughput, granular microgel method of comBO generation. **(a)** A granular microgel method was devised as an alternative to conventional bulk hydrogel approaches. **(b)** Mesodermal aggregates embedded in the granular microgel at day 5 develop vascular sprouts over the following 7 days and mature without the requirement for manual extraction (scale bar = 100 μm). **(c)** Flow cytometry at day 35 of differentiation confirms the equivalent generation of key stromal and haematopoietic lineages using the granular microgel approach. (d) Quantified relative proportions of haematopoietic and stromal cell types in each flow panel. (*n =* 2 for bulk gel, *n = 3* for granular gel. Error bars are standard deviation). **(e)** Both protocols generate a lumen-forming vascular network as confirmed by CD34 IHC in paraffin embedded sections (scale bar = 100 μm, insert = 25 μm). Confocal imaging confirms **(f)** lymphoid cells (CD56+ and CD3+ cells, scale bar = 200 μm), **(g)** osteopontin deposition (scale bar = 200 μm) and **(h)** CD34/CD44 double positive haematopoietic cells (scale bar = 100 μm, cropped regions = 25 μm). **(i)** Cytospins of dissociated comBO reveal a complex mix of morphologies annotated in detail in supp fig 12 (scale bar = 50 μm). (Step-by-step description of granular microgel protocol described in Supp Fig 9. Gating strategy for flow cytometry in Supp Fig 10. Representative plots and quantification of comBO generated from multiple iPSC lines shown in Supp Fig 11. Schematic generated on Biorender.com.)

A direct comparison of parallel differentiations performed to generate comBOs using bulk hydrogel and granular microgel methods showed no loss of cell populations or architectural complexity using the microgel approach (**Fig 2c-e**). Flow cytometry confirmed the generation of myeloid, lymphoid and stromal lineages across three independent hiPSC lines (**Fig 2 c-d, Supp Fig 10, 11**). Imaging further verified the formation of lumen-forming vessels (**Fig 2e**), similar to the architecture of organoids generated using bulk hydrogels. Lymphoid (CD3+ CD56+) were detected within the comBO volume (**Fig 2f**), as well as osteopontin-rich regions (**Fig 2g**) and CD34+ CD44+ clusters of haematopoietic cells (**Fig 2h**).

Cytospins of comBO cells derived from the granular microgels highlight a range of morphologies consistent with mature myeloid cells (megakaryocytes, neutrophils erythroid progenitor clusters, macrophages) and lymphoid progenitors (**Fig 2i, Supp Fig 12**). Together, these data confirm that a granular microgel-based approach generates comBOs of identical complexity to those produced with bulk hydrogels while offering a simplified, scalable, and automatable approach, thereby overcoming a major technical hurdle in the generation of bone marrow organoids.

### comBOs support primary multiple myeloma cells *ex vivo* and capture multi-lineage pathogenesis and remodelling

Having generated a physiologically-relevant human bone marrow model, we next explored its utility to recapitulate tumour-microenvironment interactions. Osteolineage, endothelial, adipolineage, and myeloid cells have previously been shown to be perturbed in the bone marrow microenvironment in patients with multiple myeloma, and to contribute to relapse^20,31–35^. comBO is the first human model to capture these disparate elements, offering an un-precedented opportunity to study the effects of myeloma cells on the bone marrow microenvironment.

We engrafted comBOs with CD138+ cells from cryopreserved bone marrow aspirates from three patients with myeloma to generate myeloma-comBO ‘chimeroids (MM-ComBO, **Supp Table 2**), and cultured these for two weeks before analysis by scRNA-seq. As controls, organoids that were not engrafted with adult donor cells and organoids engrafted with CD34+ cells isolated from the peripheral blood of a healthy, G-CSF mobilised adult donor were processed under identical conditions (**Fig 3a, Supp Fig 13a**).

**Figure 3:**
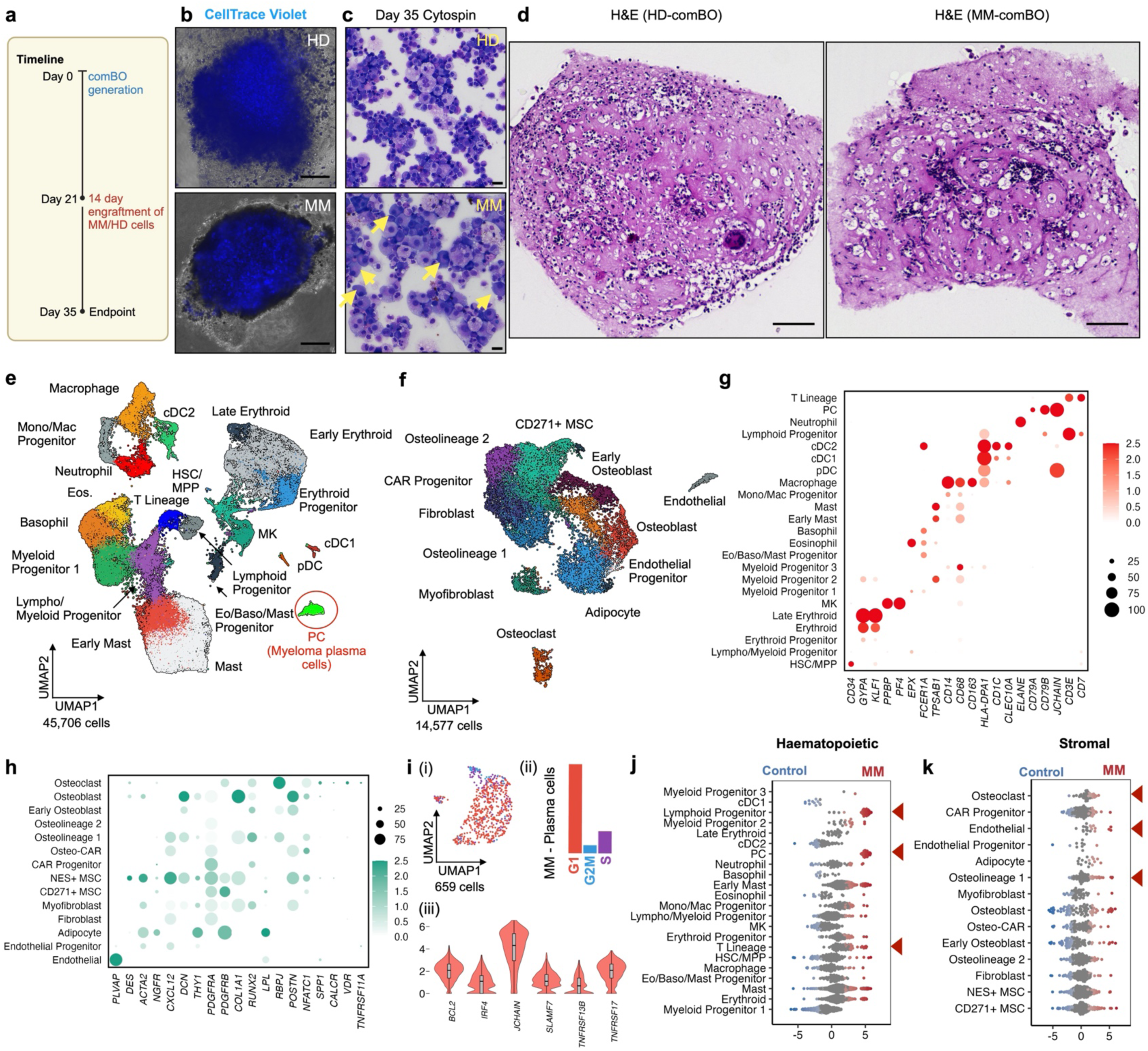
comBOs support the engraftment of primary multiple myeloma cells. **(a)** CD138+ bone marrow cells from patients with myeloma (*n* = 3) and CD34+ healthy donor (HD) cells (*n =* 1) were labelled with CellTrace Violet and engrafted into comBOs at day 21 of differentiation and cultured for a further 14 days. **(b)** Imaging of engrafted samples reveals CellTrace+ cells successfully engraft comBOs (Supp Fig 13, scale bar = 600μm). **(c)** Dissociated samples at day 14 post-engraftment (day 35 of protocol) show viable myeloma cells (scale bar = 25 μm), which are further confirmed in engrafted organoids using **(d)** paraffin embedded histological sections of control and MM-comBO (scale bar = 100 μm). **(e)** Haematopoietic cells (45,706) of the integrated myeloma, healthy donor and control un-engrafted samples with the patient specific plasma cells (PC) circled. **(f)** Stromal cells (14,577) from the same samples. **(g-h)** Simplified representative bubble plots highlighting key haematopoietic and stromal lineage markers. Full list shown in Supp Fig 14. **(i)** Myeloma plasma cells (PCs) were sub-clustered and scored using Seurat’s cell-cycle scoring method (i-ii). (iii) Cells expressing hallmark myeloma genes *(BCL2, IRF4, JCHAIN, SLAMF7, TNFRSF13B, TNFRSF17)* were observed. **(j)** Differential abundance (DA, MiloR) analysis of haematopoietic and **(k)** stromal cells highlight myeloma-induced changes in cell abundance. (Full canonical gene marker expression shown in Supp Fig 14. Schematic created on Biorender.com)

To track the adult donor cells, the healthy donor CD34+ cells and myeloma samples were labelled with CellTrace Violet on day 0 prior to engraftment, and organoids were then imaged on days 3, 7, and 14 (**Fig 3b, Supp Fig 13b**). In all engrafted organoids, fluorescent cells were observed within the organoid by day 3, and remained detectable throughout the 2-week culture period. At 14 days post-engraftment, comBOs were dissociated for scRNA-seq. Cytospins or digested organoids confirmed the presence of myeloma cells in MM-comBOs (**Fig 3c, Supp Fig 13c**). H&E staining of myeloma-engrafted comBO confirmed the presence of myeloma cells within the organoid structure (**Fig 3d, Supp Fig 13d**).

scRNAseq data was analysed and clusters annotated using canonical marker genes (**Fig 3g-h**, **Fig 1, Supp Fig 4-5**) to identify haematopoietic (45,706 cells) and stromal (14,577) cells (**Fig, 3e-f, Supp Fig 14**). Cell cycle analysis of patient-derived myeloma cells showed that, as expected, most cells were in G1, with a minority in G2M and S phase, confirming ongoing cycling but consistent with the known low proliferation rate characteristic of myeloma **(Fig 3i(i-ii), Supp Fig 13b).** These cells similarly expressed canonical myeloma markers including *BCL2, IRF4, JCHAIN, SLAMF7, TNFRSF13B, TNFRSF17* (**Fig 3i(iii)).**

Differential abundance analysis revealed the presence of lymphoid progenitors and T-lineage cells in myeloma-engrafted organoids compared to un-engrafted controls, as expected given that these organoids had been engrafted with adult donor-derived lymphoid cells (**Fig 3j**). The stromal compartment showed modest changes in the abundance of osteoclasts, endothelial and early osteoblast/osteolineage cells (**Fig 3k**). Osteoblast lineage cells demonstrate a bimodal enrichment indicating a significant shift in gene expression between myeloma and control samples, consistent with known changes to osteoblast differentiation induced in myeloma^32,34,35^.

GSEA analysis comparing myeloma engrafted comBO to controls identified significantly enriched hallmark gene sets across MM-comBOs, consistent with myeloma-induced remodelling of the comBOs (**Fig 4a**). A further analysis of the four cell types with the highest numbers of significantly enriched gene sets (CD271+ MSC, NES+ MSC, osteolineage 1, and lymphoid progenitors) highlighted inflammatory and metabolic gene sets (**Fig 4b, Supp Fig 15**). Stromal cells (CD271+ MSC, NES+ MSC, osteolineage 1) demonstrated TNF, IL6, and interferon-mediated inflammatory responses, as well as an increase in fatty acid metabolism and PPARA signalling gene sets ^20,21,31–33,36^.

**Figure 4:**
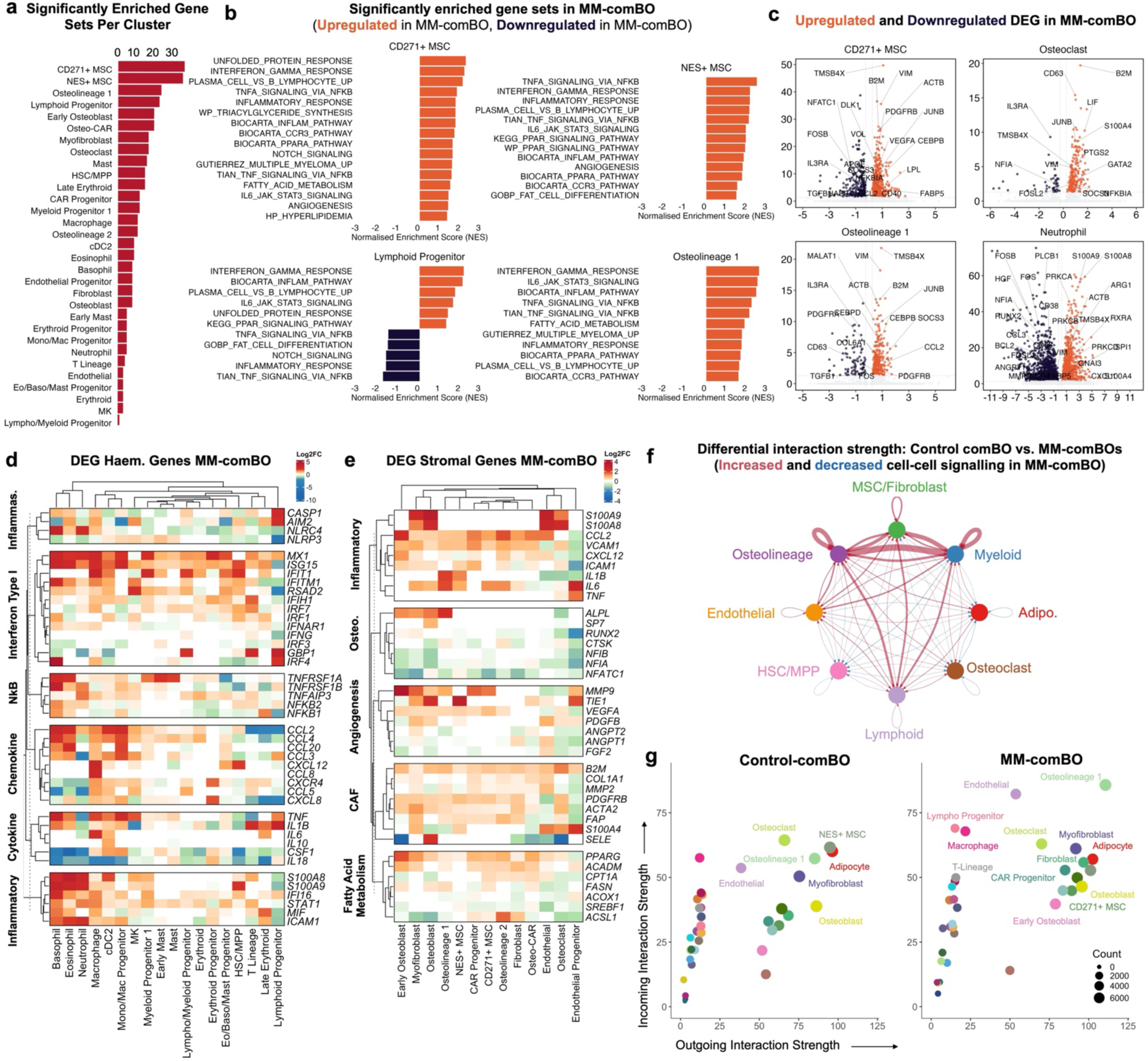
Multiple myeloma engrafted comBOs capture hallmarks of complex disease pathology. **(a)** Gene set enrichment analysis using hallmark gene sets and comparing myeloma engrafted MM-comBO with control organoids reveals an extensive dysregulation, most evident in MSC, osteolineage cells and lymphoid progenitors in MM-comBOs. **(b)** Representative GSEA plots highlight inflammatory response, fatty acid metabolism, and unfolded protein response genes as some of the major upregulated pathways in key clusters. **(c)** Volcano plots identifying key genes involved in inflammation, osteo/adipo-lineage fate, and fatty acid metabolism in key myeloma associated cell types. Heatmap of DEGs in specific pathways highlighted by GSEA in **(d)** haematopoietic and **(e)** stromal cells. **(f)** CellChat analysis identifies a dysregulation of cell-cell interactions across key clusters, identifying upregulated interactions in MM-comBO. **(g)** Analysis of net outgoing and ingoing signals reveals a substantial change in outgoing cell signals in MM-comBO, consistent with a remodelling of the organoid because of engraftment.

Volcano plots of key populations demonstrated an upregulation of key DEGs in cell types identified as dysregulated in GSEA (**Fig 4b**-**c**). Stromal activation markers (*ACTB, PDGRB, VIM),* angiogenic factors *(VEGFA*), adipogenic genes *(LPL, FABP5, APOE),* and inflammatory genes *(NFKBIA, CCL2)* are significantly up-regulated in CD271+ MSC (**Fig 4c**). Osteoclasts demonstrate an upregulation in markers supporting osteoclast differentiation (*PTGS2, JUNB, FOSL2)* and indicating osteoclast activation and inflammation (*LIF, S100A4, NFKBIA, B2M)* (**Fig 4c**)^31,34^. Early osteolineage cells (osteolineage 1) demonstrate an upregulation of *JUNB*, induced by inflammation and indicating impaired osteogenesis (**Fig 4c**). Similarly, an upregulation of *SOCS3* is a consequence of IL6 mediated inflammation, known to suppress osteoblast differentiation. *CCL2, CEBPB, PDGFRB* up-regulation similarly indicate an activated stromal compartment, with *CCL2* known to drive the recruitment of monocytes/macrophages that can facilitate osteoclast differentiation in myeloma^34^.

A volcano plot of neutrophils, recently reported as a major driver of myeloma inflammation^21^, demonstrate a significant upregulation of inflammatory markers including *S100A8/A9*, alarmin axis proteins now known to impede therapeutic efficacy in myeloma^37^.

Heatmaps of Log2FC in significantly differentially expressed genes confirmed an inflammatory and metabolic dysregulation of MM-comBO compared to un-engrafted and healthy donor engrafted organoids (**Fig 4d-e**). An upregulation of interferon type I response genes, and NFkB mediated TNF-α signalling were observed across the Eo/Baso/Mast arm, as well as in dendritic cells (cDC2) and mono/mac lineage cells (**Fig 4d**). A similar upregulation in chemokines (e.g. *CCL2, CCL3, CCL4*) and alarmin axis (*S100A8, S100A9*) mediated inflammation was also observed in neutrophils and Eo/Baso/Mast cells^8,20,21,37^. Inflammasome related *NLRC4* was upregulated in neutrophils and basophils, while *AIM2* and *CASP1* were upregulated in lymphoid progenitors in the MM-comBO.

In the stromal compartment, inflammatory genes (*S100A8, S100A9, CCL2, VCAM1, IL6, TNF*), differential expression of osteolineage gene sets (e.g. the osteoblastic maturation genes *NFIA*, and *NIFIB)*, and an increase in angiogenesis genes (*TIE1, VEGFA, ANGPT2*) were observed (**Fig 4e**). Similarly, genes associated with fibroblast activation in cancers (e.g. *FAP, S100A4, MMP2, COL1A1, ACTA2*) were significantly upregulated, and dysregulation of fatty acid metabolism was observed (*PPARG, ACADM*, and *CPT1A)* (**Fig 4e**). Together, GSEA and DEG analysis show that the engrafted malignant plasma cells extensively remodel the comBO microenvironment, recapitulating multilineage hallmarks of myeloma pathogenesis.

### Cell-cell interaction analysis reveals a druggable MIF-mediated inflammatory pathway

To investigate dysregulated cell-cell crosstalk in MM-comBOs, CellChat analysis was performed^38,39^. The resulting interactome identified significantly upregulated interactions in MM-comBOs, particularly in predicted ligand-receptor interactions across osteolineage cells, MSC/fibroblasts, consistent with DEG and GSEA data (**Fig 4f, Supp Fig 16**).

Analysis of incoming and outgoing interaction strength, defined as total receptor and ligand expression respectively, demonstrated a significantly altered distribution in MM-comBO samples when compared to controls. In MM-comBO (osteolineage 1, myofibroblast, adipocytes, endothelial cells, CAR progenitors and osteoblasts) demonstrating relatively increased outgoing signals. Conversely, lymphoid progenitors, T-lineage cells, macrophages, and endothelial cells exhibited the most marked shift in incoming interaction strength (**Fig 4g**). Combined with GSEA and DEG analysis showcasing myeloma induced dysregulation, including significant inflammation, this data illustrates a significant, multi-lineage remodelling of comBO in response to infiltrating myeloma cells. Together, this comparative data set identifies a significant dysregulation of cell-cell interaction induced by myeloma cells within comBOs.

An analysis of receptor-ligand interactions mediated by myeloma cells (PC) within MM-comBO identified distinct pathways that differentially target stromal and haematopoietic cells (**Fig 5a**). *MIF-(CD74+CD44*), *MIF-(CD74-CXCR4)* and *CCL3- CCR1* mediated interactions were the highest confidence pathways between MM cells and dendritic cells, lymphoid progenitors, and macrophages. In contrast, *SPP1-* mediated interactions were the strongest interacting partners between MM and stromal compartment. Expression of *MIF, SPP1*, and *CCL3* was evident in myeloma cells (**Fig 5b**), although *TNF* was not expressed, suggesting that the TNF-α signalling enriched in MM-comBOs was due to secondary effects – i.e. pro-inflammatory reprogramming of myeloid cells in the myeloma microenvironment. DEG analysis identified (**Fig 4d**) myeloid cells (macrophage, cDC2, basophils, eosinophils) as the major source of TNF-α in MM-comBO.

**Figure 5:**
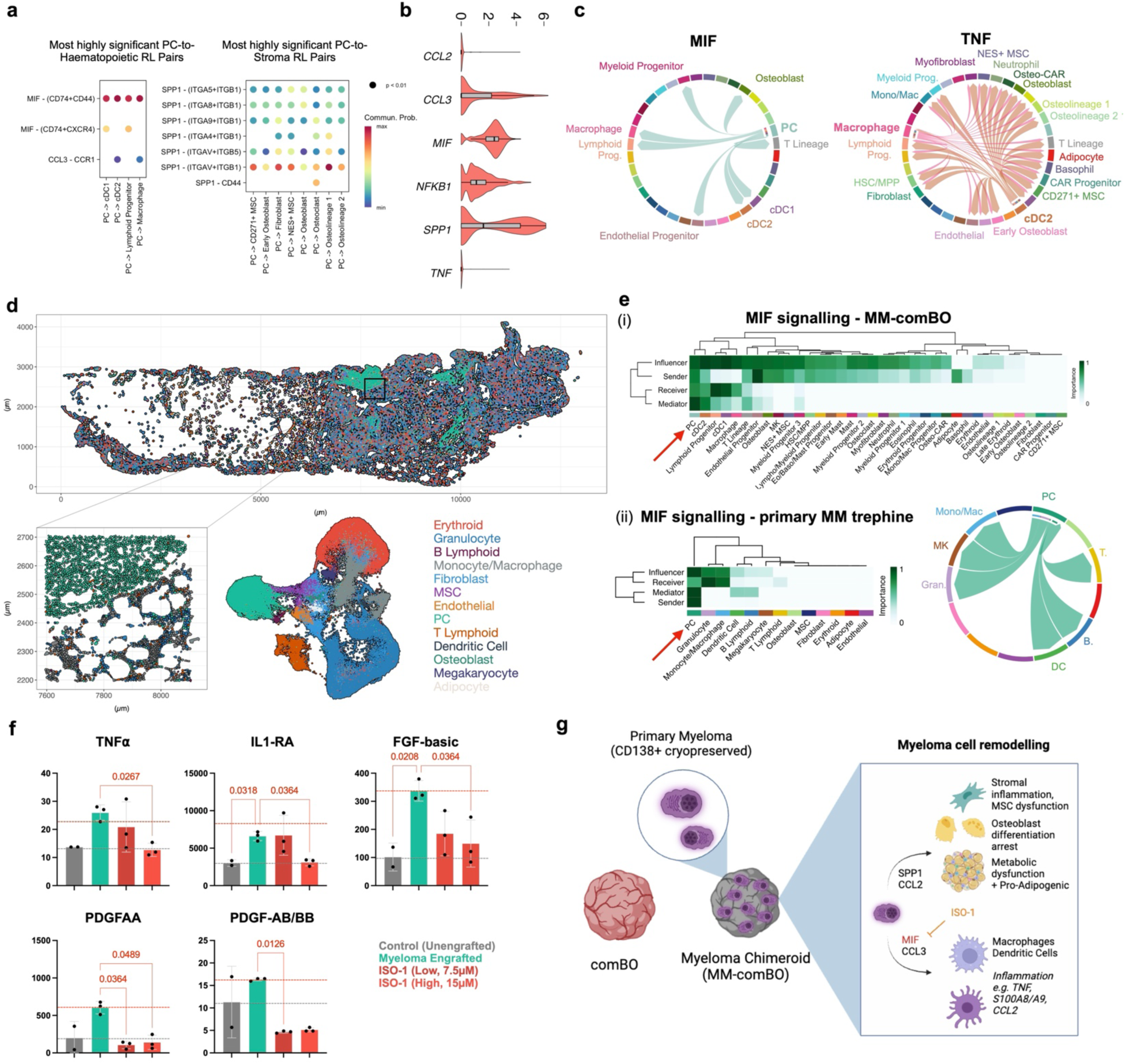
Cell-cell interaction analysis reveals mechanisms underpinning niche remodelling in myeloma potential therapeutic avenues. **(a)** CellChat analysis reveals MIF, CCL3, and SPP1 as major ligand-receptor (LR) interactions originating from myeloma plasma cells (PCs), with MIF interactions highlighted as the most significant. **(b)** Violin plots confirm the expression of *MIF, SPP1*, *and CCL3*, however while *TNF* is one of the major pathways disrupted in MM-engrafted comBOs, this is not expressed in patient cells despite its upregulation in myeloid cells (Neutrophils, DCs, Eo/Baso/Mast cells) in MM-comBOs. **(c)** Chord diagrams identify and confirm myeloma plasma cell-derived MIF signalling to cDC2 and macrophage clusters, which in turn are responsible for the inflammatory TNF signals observed. (Chord diagrams depicting source of signalling pathway and colour coded by cluster) **(d)** Xenium analysis of primary patient trephines captures a range of haematopoietic and stromal cell types. **(e)** (i) A clustered heatmap of centrality scores (CellChat) identifies plasma cells as the major source of MIF-mediated effects in the comBOs, confirmed by (ii) primary Xenium analysis of patient bone marrow trephine. **(f)** MIF inhibition was tested using patient engrafted MM-comBOs (Full figure in Supp Fig 14). Samples from 3 patients were engrafted for 5 days before treatment with ISO-1 (MIF inhibitor) at low (7.5 μM) and high (15 μM) doses. Un-engrafted (gray) controls were taken from un-engrafted samples from two separate differentiations (n = 2). For myeloma engrafted samples, each individual data point represents supernatant pooled from 8 organoids engrafted with cells from a single patient. MIF inhibition (ISO-1 treatment) significantly mitigated the upregulation of TNFα, IL1Ra, FGF-basic, PDGF-AA and PDGF-AB/BB (Error bars are standard deviation. Significance determined with a Kruskal-Wallis test (Uncorrected Dunn’s test, alpha threshold and confidence level = 0.05). **(g)** Together, this data suggests differential interactions that drive elements of the MM mediated pathological remodelling of comBOs, capturing hallmarks of disease.

Mapping the MIF signalling interactome confirmed that plasma cell-derived MIF was received primarily by myeloid cells, including dendritic cells and macrophages (**Fig 5c**). Up-regulated TNF signalling in MM-comBO was in turn primarily derived from macrophages and dendritic cells (cDC2) (**Fig 5c**). Together, this data suggested that plasma cell-derived MIF underpins TNF-mediated inflammation in MM-comBOs.

We looked to confirm the relevance of these findings in clinical specimens. Xenium spatial transcriptomics using a custom panel was applied to bone marrow trephines from patients with newly diagnosed and relapsed disease (n= 3). Xenium data was annotated to identify plasma cells, haematopoietic and stromal cells (**Fig 5d**). CellChat network centrality analysis was performed on both primary trephine data (Xenium) and MM-comBOs (**Fig 5e, Supp Fig 17**). In both samples, clustered heatmaps of critical senders (ligand expressing cells), receivers (receptor expressing cells), mediators (expression of both receptors and ligands) and influencers (combination of sending, receiving, and mediating roles) in MIF signalling identify plasma cells as the critical driver of MIF signalling networks (**Fig 5e**, **Supp Fig 17**).

To functionally validate the role of MIF in myeloma-mediated inflammation, we designed inhibition experiments. MM-comBOs were generated using MM cells from 3 patients, and a MIF inhibitor (ISO-1) was introduced at day 5 post-engraftment. At day 14 post-engraftment, supernatant was taken from treated samples and subject to luminex analysis for markers of inflammation and metabolic dysfunction.

Inflammatory markers (TNF-α, IL1-RA) and stroma mediating factors (FGF-basic, PDGF-AA, and PDGF-AB/BB) were upregulated in myeloma-engrafted organoids (**Fig 5f**). The MIF inhibitor ISO-1 significantly decreased this up-regulation (**Fig 5f**), indicating an effective inhibition of MM-induced niche inflammation and stromal/endothelial interactions.

Together, our data report an extensive remodelling of comBO induced by myeloma cell engrafting, re-capitulating hallmarks of disease (inflammation, transcriptional re-wiring of MSCs and osteolineage cells). This data highlights potential mechanisms by which myeloma plasma cells drive remodelling, including a MIF signalling axis which was validated in primary patient trephines and through inhibition assays (**Fig 5g**).

## Discussion

In this work, we present comBO, a multi-niche organoid system that includes the cellular elements required to model the osteoblastic, lymphopoietic and myeloid compartments of human bone marrow. Unlike previous bone marrow organoid platforms that model the myelopoietic niche^16–18^, comBO captures both the myeloid micro-environment of the central marrow and the osteoblastic niche critical for lymphopoiesis, HSC maintenance, and modelling diseases of the haematopoietic tissue^3,5,7,41^. By combining physiologically relevant oxygen levels with graded cytokine exposure and a 3D scaffold, comBO successfully integrates these distinct niches to more faithfully capture the complexity of the native tissue.

Balancing complexity with scalability and reproducibility has been a persistent challenge in organoid research. The microgel-based approach used in comBO overcomes limitations of bulk hydrogel systems, enabling a single-step differentiation protocol that is cost-effective, reproducible, and scalable. This protocol involves a significant reduction in hands on time by removing labour intensive extraction (30-60 minutes per 96-well plates) and sequential gel polymerisation steps (2 hours per step), and a similar total reduction in hydrogel materials (Matrigel, collagens, fibrinogen) of 80%. This is a critical step towards implementing highly complex models in high-throughput discovery and validation pipelines.

Multiple myeloma was chosen as a proof-of-concept application because it remodels its bone marrow microenvironment to support growth and modulate therapy response ^8,20,31,32,36,42–44^. Primary myeloma cells have classically been extremely difficult to culture *ex vivo*, and mouse models poorly emulate the disease, major limitations in both fundamental and translational research^42^. Moreover, no culture platform to date has captured the diverse stromal and haematopoietic cellular elements that contribute to disease progression, relapse and the characteristic disruption of bone homeostasis of multiple myeloma. ComBO-engrafted primary myeloma cells survive and cycle, recapitulating complex interactions with the microenvironment and providing a platform to dissect mechanisms underpinning MM pathogenesis and drug responses.

Inflammation is known to play a crucial role in myeloma niche remodelling. Pro-inflammatory cytokines such as IL6, TNFɑ, IL1B, CCL, IL1-RA, and complement factor, and PDGF-AA have been reported as markers of inflammation in myeloma infiltrated bone marrow, contributing to bone disease by enhancing osteoclast activation and inhibiting osteoblast function^8,20,21,45^. Targeting inflammation in myeloma holds therapeutic relevance in preventing the progression of precursor disease, bone loss and enhancing responses to immunotherapy, making it a key area of priority research.

Our data implicate MIF in MM-mediated inflammation, a finding consistent with reports linking MIF to T-cell exhaustion and resistance to proteasome inhibitors in MM^40,46^. Using spatial transcriptomic data from patient trephines and pharmacological inhibitors, we validated the role of MIF and demonstrated its potential as a therapeutic target. MIF inhibition through ISO-1 significantly decreases secreted TNFɑ, IL1-RA, FGF-basic, PDGF-AA, and PDGF-AB/BB in comBOs engrafted with myeloma cells from 3 different patients.

This data highlights the utility of comBO as a pre-clinical platform that both identifies and validates a potential target using patient cells. In the broader context of bone marrow biology and disease, inflammation and stromal re-programming are known contributors to disease progression which, in the absence of complex models, have thus far been difficult to study *ex vivo*^7,8^.

Future studies will explore the dynamics of myeloma-driven inflammation and stromal re-modelling, with a focus on MIF inhibition and its effect on myeloma cell survival and proliferation in combination with plasma cell targeting therapies. Additionally, validation of comBO’s ability to support primary human HSCs *ex vivo* via the osteoblastic niche could further establish its utility for translational research. In the absence of dynamic flow, the vasculature generated by comBO and other static organoid systems is not long-lived. Implementing microfluidic strategies to perfuse the vasculature will be a major focus of future work, enabling long-term cultures and thus the assessment of clonal evolution in blood cancers and mechanisms of acquired resistance to therapy.

In summary, comBO represents a significant advancement in organoid technology, providing a scalable and physiologically relevant platform to study human haematopoiesis and bone marrow pathologies. By modelling multiple bone marrow niches, comBO bridges critical translational gaps and enables mechanistic studies of blood disorders. comBO represents one of the most faithful human organoid models in terms of cell complexity and function, showcasing the potential and power of these systems as tools for discovery and translational research.

## Materials and Methods

### Cell culture

Human induced pluripotent stem cells were maintained on Geltrex (ThermoFisher Scientific, Cat#A1569601) coated plates in StemFlex (ThermoFisher Scientific, Cat#A3349401) maintenance medium. Three hiPSC lines were used in this study, Gibco hiPSC (ThermoFisher, Cat#A18945), SCTi003-A (Stem Cell Technologies, Cat# 200-0511), and KOLF2.1J (The Jackson Laboratory, Cat# JIPSC1000). Cells were maintained during routine passaging in a 37°C incubator at 5% CO_2_, with routine clump passaging performed using ReLeSR (Stem Cell Technologies, Cat#100-0483). Cells were mycoplasma tested at 1-month intervals, with karyotyping performed on purchase/import.

### Organoid differentiation

A full list of all media compositions and cytokine cocktails is provided in **Supp Fig 1**. Briefly, the comBO differentiation protocol involves a clump dissociation (day -1) in StemFlex supplemented with RevitaCell (ThermoFisher Scientific, Cat#A2644501) and subsequent culture in ultra-low attachment 6-well plates. On day 0, the resulting aggregates are collected by centrifugation (100Gx 3 minutes, room temperature). Media is then aspirated before cells were resuspended in APEL2 (Stem Cell Technologies, Cat#05275) supplemented with BMP4, FGF2, VEGFA and CHIR99021 (Supp Fig 1) and cultured in 5% oxygen. Note that cells are maintained in 5% O2 for a total of 20 days under the comBO protocol, and as such care must be taken when performing media changes to avoid a prolonged exposure to atmospheric oxygen. On day 3, embryoid bodies are collected by gravitation and media is then aspirated before resuspension in APEL2 supplemented with VEGFA, BMP4, FGF2, hSCF, FLT3L (Supp Fig 1). On day 5, cells were either prepared for embedding in bulk hydrogels (data from Fig 1-3), or in a granular microgel. Bulk hydrogel samples were prepared as follows: First, a 12-well tissue culture plate was washed with 1mL phosphate buffered saline (PBS). For each well required, 500 µL of hydrogel is prepared.

The first (base layer) is a fibrin-collagen type-1 mixed hydrogel generated by mixing fibrinogen (Cat# F8630-5G, Merck) and VitroCol (Advanced Biomatrix, Cat #5007), with a hydrogel diluent, thrombin (2 U/mL), and 1M NaOH on ice to a final concentration of 1.25 mg/mL and 2 mg/mL respectively. The hydrogel diluent described by Olijnik *et al.* is used to dilute the hydrogel mixture. Once prepared and added to a 12-well tissue culture plate, the first layer of hydrogel is incubated at 37°C for 90 minutes. Once the base layer is polymerised, cell aggregates are collected by gravitation and resuspended in the second hydrogel layer. Approximately 40 aggregates are cultured in each well. The second layer is comprised of fibrinogen (1.25 mg/mL), thrombin (2 U/ml), VitroCol (2 mg/mL), and collagens type:I-IV (3 mg/mL, 1 mg/mL respectively, diluted with hydrogel diluent, and neutralised with NaOH).

Note that thrombin and NaOH should be added as the last components of the hydrogel mix, which should be mixed and maintained on ice. For a further, detailed step-by-step protocol for hydrogel preparation, please refer to Olijnik *et al.* Once mixed, the cell-containing second hydrogel layer should be added dropwise to the first gel layer before incubation at 37°C, 5% CO_2_, for 90 minutes. Once fully polymerised, prepare 1 mL of APEL2 for each well of a 12-well plate as described in Supp Fig 1 and add dropwise. On day 7, 1 mL of media is added as described in Supp Fig 1. From day 10 onwards, media changes are performed at a ratio of 50:50 conditioned to fresh medium. On day 12, sprouted aggregates are collected using a pasteur pipette and sub-cultured in 96-well ULA plates as individual organoids, with media changes as described in Supp Fig 1.

For granular microgels, a volume of layer 2 hydrogel solution is prepared on day 5 and transferred into a sealed sterile syringe and allowed to polymerise for 90 minutes before fragmentation by passing through a series of sterile needles (18G, 21G, 23G, 25G, 27G). The resulting fragmented gel is resuspended in a volume of APEL2 day 5 medium (Supp Fig 1). A total of 450µL of gel is prepared per 96-well plate required, diluted to a final volume of 5 mL in day 5 medium. A total of 120 cell aggregates are re-suspended in a 5 mL volume to ensure a complete 96-well plate. The final 5 mL volume, comprised of cell aggregates, growth factors, and fragmented microgel, is distributed into a 96-well ULA using a reagent reservoir, with a volume of 50 µL per well dispensed. On day 7, a further 50 µL of complete media is added per well. From day 10 onwards, partial (50:50) media changes are performed using a multi-channelpipette. Note that users should take care not to accidentally aspirate organoids at early stages, a schematic detailing this protocol is shown in **Supp Fig 9**. For troubleshooting and establishing protocols, we would recommend readers join our online community (https://groups.google.com/g/morerganoids) in which many FAQs and experimental details have been discussed.

### Flow cytometry

Organoids were collected by gravitation into tubes and washed twice with PBS prior to dissociation. Dissociation was performed using collagenase D (Roche, Cat#11088866001, dissolved in HBSS (Sigma, Cat#H9394)) at a final concentration of 2.5 mg/mL and supplemented with 1% FBS and 1000 U/mL DNase I in IMDM medium, as described by Schreurs *et al*. Typically, 1 mL of this dissociation solution could be used for dissociating 20 organoids. Organoids were incubated at 37°C for 15-30 minutes, with pipette trituration performed every 5-8 minutes, until a single-cell suspension was obtained. Digestion was stopped by adding a stopping buffer consisting of PBS, 10% FBS, and 20 mM EDTA. Cells were then centrifuged at 300g for 5 minutes, and the cell pellets were resuspended in flow cytometry buffer (PBS + 10% FBS + 2 mM EDTA) for antibody staining. Following staining, samples were washed with flow cytometry buffer and analysed using a BD LSR Fortessa X20 flow cytometer. Single-stained controls and fluorescence-minus-one controls were used for the panel setup with antibodies listed in Supp Table. 1. Flow cytometry data were analysed in FlowJo v.10.10 software.

### Imaging

Organoids used for imaging were fixed with 4% paraformaldehyde on a roller at room temperature for 1h, then stored in PBS at 4°C for further processing. Imaging of fluorescently labelled organoids (Antibody details in Supplementary Table 3) was carried out using a Zeiss LSM 880 Axio Observer Z1 confocal microscope, running Zen 2.3 SP1 FP3 (Black), with a Plan-Apochromat 10x/0.45 or a C-ACHROPLAN 32x/0,85 W Korr VIS-IR objective at 0.6 x zoom, taking Z-stacks of various height and scaling [1.384 x1.384x8µm @10x and 0.216x0.216x3µm @32x], with a pixel dwell time of 0.7 µs and 4 x line average. Fluorescent labelling and subsequent clearing with ethyl cinnamate was performed as described by Olijnik *et al* ^17^. Histology imaging was performed using an EVOS M5000 (Thermo Scientific). Histology imaging was white balanced using a custom ImageJ macro. Fluoerscent images were pre-processed using background subtraction (rolling ball) before LUT adjustment for visualisation purposes. Typically, 3D projections were used for Z-stack images.

### Single cell RNA sequencing

For all single-cell RNA sequencing experiments, cells were from a total of 4 independent differentiations. For day 20 comBO sequencing, these were cryopreserved and thawed before live cell sorting and processing with the Chromium Next-GEM 3’ kits v3.1, 4 rxns (Cat#1000269, 10X Genomics) using the Dual Index 3’ protocol (CG000315). For day 35 comBO, healthy donor and MM patient-engrafted samples were prepared fresh on day 35 of the experiment by collagenase D dissociation and live cell sorting before loading on the 10x Genomics 3’ HighThroughput kit v3.1 (Cat#1000370, 10X Genomics). 2 unengrafted controls were used for this experiment, each from independent differentiations. One was cryopreserved and thawed prior to live cell sorting, the other was prepared in parallel to engrafted samples.

### Analysis of single cell RNA sequencing

Single cell data was processed using the CellRanger pipeline (v. 7.0.0), post-processed using CellBender (v.0.3.0)^47^, Souporcell (v.3.0.0)^48^Seurat (v.5.1.0) ^49,50^.^48^. Predicted ligand receptor (LR) analysis was performed using CellChat (v.1.6.1)^38,39^. Differential abundances were computed with miloR package (v.2.0)^51^, integration performed using Harmony (v.1.2.0) ^52^ and plotting performed using the scCustomise package (v.2.1.2). All raw data and processed data (R objects) are available to review on GEO accession GSE287648: https://www.ncbi.nlm.nih.gov/geo/query/acc.cgi?acc=GSE287648. All analysis scripts are available via GitHub (https://github.com/khanaswimm/comBO.v1. These repositories are private until after peer review and publication, at which point they will become publicly available.) In summary, all CellRanger-aligned cells were processed for ambient RNA removal via CellBender, patient demultiplexed via Souporcell and imported as Seurat objects. Quality control filtering was performed on RNA features, counts, and percentage mitochondrial genes before normalization via SCTransformation (genes regressed were mitochondrial, ribosomal,and cell cycle genes). Typically, QC standards were: nFeature_RNA (min. 500, max 10,000), nCount_RNA (min. 1000, max. 60,000), and percentage mitochondrial genes below 15%. This would vary depending on data set, but is fully documented in the relevant GitHub.

Data was then integrated with the Harmony package to form batch corrected seurat objects. (integration co-variance was sample) Based on the expression of canonical haematopoietic and stromal genes, objects were divided into haematopoietic and stromal objects for cluster analysis.

LR CellChat analysis was performed on all cells, merging the separated haematopoietic and stromal compartments. To highlight broad changes in LR interactions CellChat analysis was first performed on merged data from all of the organoid studies (either mBMO and d20 comBO, or control and MM-comBO data). Further analysis focussed on identifying key condition-specific LR pairs and changes on either d20_comBO data or MM-comBO data.

Differential Gene Expression and Gene Set Enrichment analysis were performed using Seurat’s built in find markers function and the fGSEA package. Where a cluster was only present in one of the data sets, this cluster would be removed to facilitate comparative analysis. Typically, an FDR of 0.20 was used with a logfc threshold based on the data in question (average 0.25). GSEA analysis was performed using a hallmark gene set which included bone marrow and inflammatory gene sets (uploaded in relevant Github). GSEA and DEG were performed by comparing like-for-like clusters in control (un-engrafted and HD engrafted) vs. MM engrafted organoids. Differential Abundance (DA) analysis was performed comparing un-engrafted controls vs. MM-comBO. Finally,

### Methocult cultures

Live CD45^+^CD34^+^CD144 cells were sorted from day 21 dissociated comBOs on a FACSAria Fusion sorter (Becton Dickinson). Cells were plated in methylcellulose H44335 Enriched (Cell Technologies). After 14 days, colonies derived from erythroid (BFU-E, burst forming unit-erythroid) and granulo-monocytic (CFU-GM, colony forming unit granulocyte macrophage) progenitors were counted.

### Patient cell engraftments

Healthy donor CD34+ cells were obtained through the INForMed Study, University of Oxford (IRAS: 199833; REC 16/LO/1376). Written informed consent was received for all the participants for the donation of human samples. Primary cells used for myeloma comBO engraftment, and bone marrow trephine samples used in spatial transcriptomic experiments, were all obtained from the Oxford Radcliffe Biobank, REC approval reference 19/SC/0173.

### Inhibitor experiments

Inhibitor experiments were performed by engrafting primary myeloma samples on day 21 of comBO differentiation. Cells from three myeloma patients were seeded at 20,000 cells per organoid and cultured for a further total of 14 days. Inhibitors were added at day 5 post-engraftment once cells had engrafted and recovered from thaw and engraftment. All inhibitors were added in a 50 µL volume at 2x concentration (with 50 µL being the volume within each organoid well prior to treatment). The final volume per well was 100 µL, and concentrations are listed as final concentration values. ISO-1 (MIF inhibitor) was used at 7.5 µM (low dose) and 15 µM (high dose), with re-dosing (re-feeding) every 72 hours (Cat# HY-16692, MedChemExpress). At day 9 after treatment (14 days post-engraftment), supernatant from 8 organoid wells under the same condition was pooled and subject to Luminex cytokine assays.

### Statistical analysis

Statistical analysis performed on inhibitor experiments was a Kruskal-Wallis with multiple comparisons to myeloma engrafted samples (Uncorrected Dunn’s test, alpha threshold and confidence level = 0.05). This analysis was performed using GraphPad PRISM 10 (v. 10.3.1).

### Luminex multiplex analysis

For Luminex analysis, supernatant from 8 organoids per treatment condition and patient were pooled and spin at 500xG for 5 minutes before being placed on dry ice and subsequently stored at −80°C. Protein expression was measured using the Human XL Cytokine Luminex Kit performance Assay (Cat# FCSTM18B, R&D Systems). Analysis was restricted to factors upregulated in engrafted samples when compared to un-engrafted controls.

### Xenium spatial transcriptomics

Bone marrow trephine specimens were fixed in formalin, decalcified and then paraffin embedded. Specimen blocks were sectioned and mounted on Xenium slides, then underwent deparaffinisation and decrosslinking according to the manufacturer’s protocol. Targeted probes designed for a custom 480-parameter panel were applied along with cell segmentation staining according to the Xenium v1 workflow (protocol CG000749, revision A). Data were then acquired with the Xenium Analyzer. Downstream analysis was performed in R, using Seurat and CellChat.

### Code availability

No new code was generated for this manuscript, all analysis scripts are available via GitHub: All analysis scripts are available via GitHub (https://github.com/khanaswimm/comBO.v1). Repository is private until peer review and publication, at which point this and all other raw data will be made publicly available.

### Data availability

All raw data and processed data (fully annotated R objects) are available to review on GEO accession GSE287648: https://www.ncbi.nlm.nih.gov/geo/query/acc.cgi?acc=GSE287648.

Repository is private until peer review and publication, at which point this and all other raw data will be made publicly available.

### Contributions

AOK, YS, and JSR performed cell culture experiments (hiPSC maintenance, organoid differentiation, drug dosing, and patient engraftments). YS performed flow cytometry experiments, YS and CB performed formal analysis of flow cytometry data. AOK, YS, SK, CGS, APC assisted in preparing sample for single cell RNA sequencing. AOK, ARR, KG performed analysis of single cell RNA sequencing. JR performed Luminex assays. SG and EW prepared myeloma samples, EW performed formal analysis and preparation of Xenium samples. RH and SAA performed histological and morphological analysis. AOK, SG, BP and YS co-authored manuscript. AOK conceived of the study, and AOK and BP sought funding for the project. All authors reviewed data and the manuscript in its preparation.

## Supporting information

Shen et al. Supplementary Material

## Acknowledgements

The authors would like to thank the exceptional core facilities at the MRC WIMM for their support and guidance (Jana Koth and Cyril Lai from the Wolfson Imaging Centre, Sally-Ann Clark *et al.* Flow Cytometry Core, and Maria Greco *et al.* Sequencing Core (Advanced Single Cell Omics Facility). The authors would also like to thank patients for their participation, and Prof. Xin Lu and Dr. Xu Liu for their generous provision of the KOLF2.1J iPSC line. We would like to acknowledge the National Institute for Health Research (NIHR) and Oxford Biomedical Research Centre (BRC).

