## Supplementary material for "comBO: A combined human bone and lympho-myeloid bone marrow organoid for pre-clinical modelling of haematopoietic disorders": Shen et al. Supplementary Material

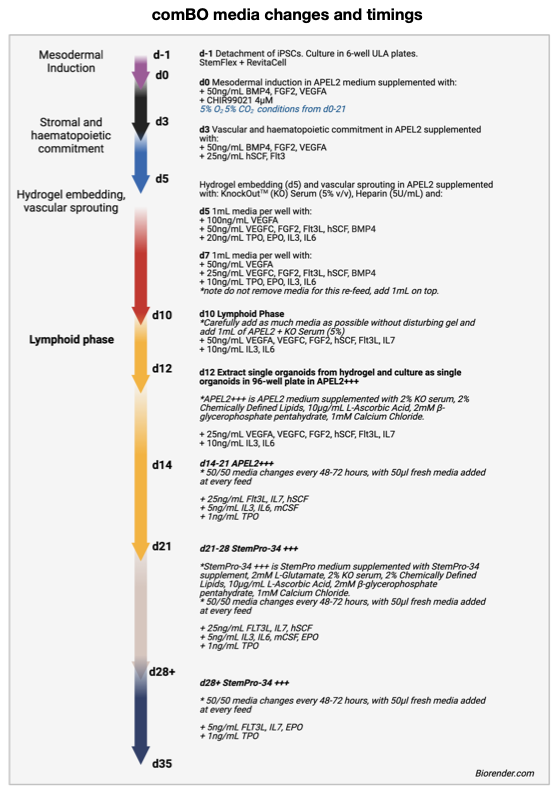
**Supplementary Figure 1:** Full list of media changes, compositions, and timings for the comBO protocol. Schematic created on [**biorender.com.**](http://biorender.com)

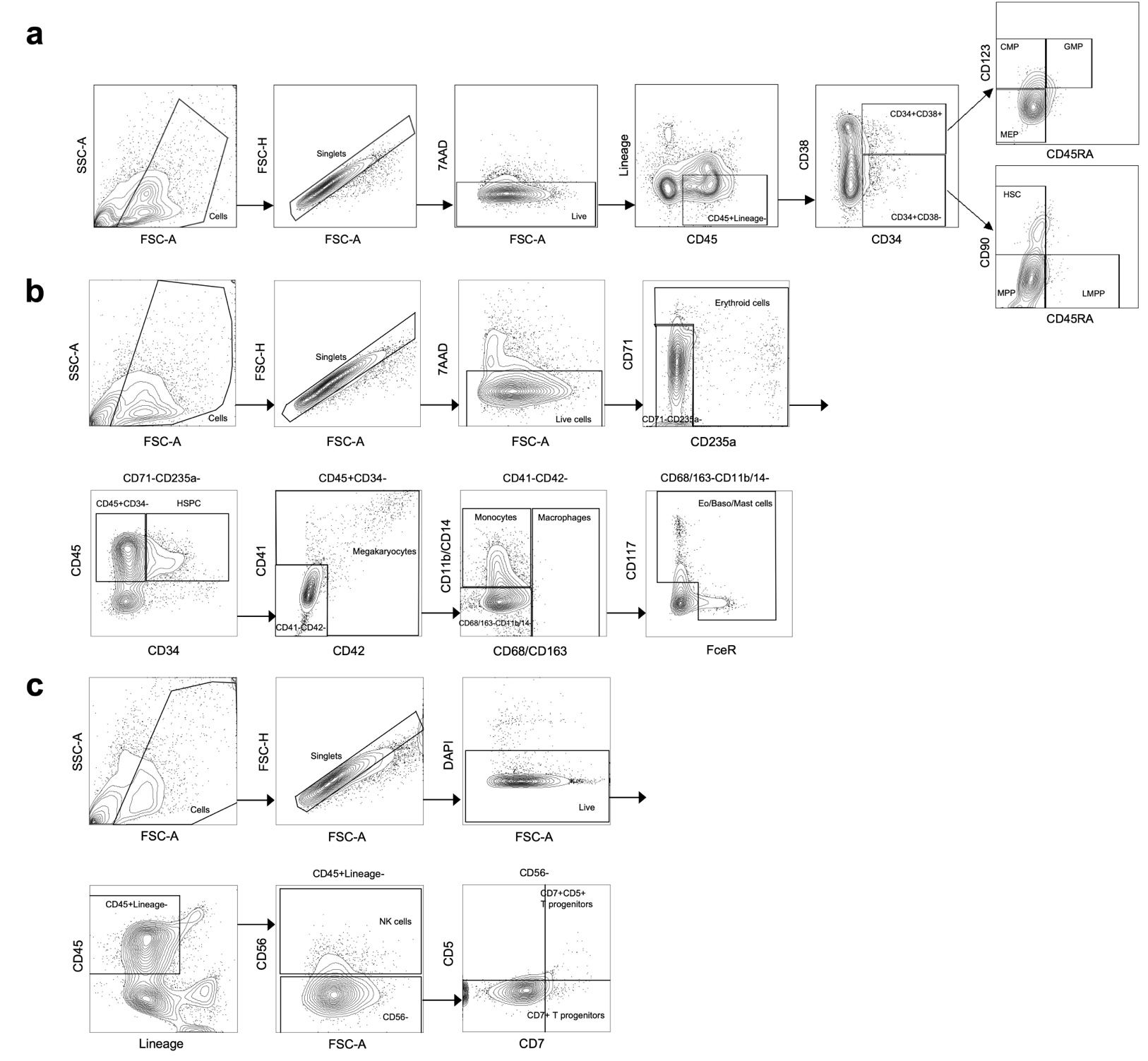

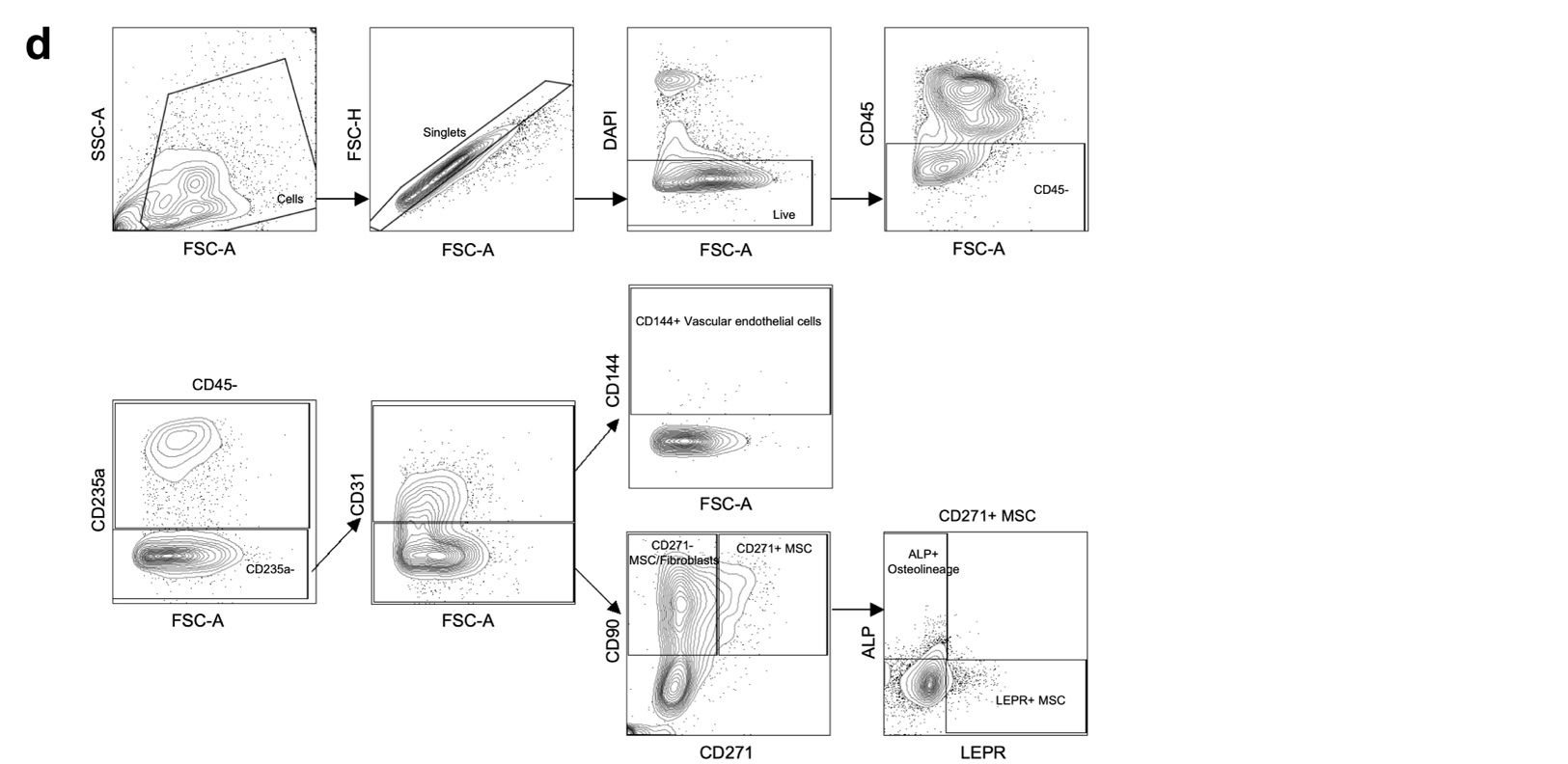

**Supplementary Figure 2:** Gating strategy for flow cytometry data reported in Fig. 1a-b. (a) Gating strategy for HSPC panel. CD45+ lineage- haematopoietic cells are gated from live singlets. Then CD34+ CD38- cells are classified as HSCs (CD90+ CD45RA-), MPPs (CD90- CD45RA-), or LMPPs (CD90- CD45RA+) based on their expression of CD90 and CD45RA. CD34+ CD38+ cells are classified as CMPs (CD123+ CD45RA-), MEPs (CD123- CD45RA-), or GMPs (CD123+ CD45RA+) based on their expression of CD123 and CD45RA. (b) Gating strategy for myeloid panel. After gating live singlets, erythroid cells are identified as CD71hi CD235hi. Then CD45+ CD34+ cells are classified as HSPCs. From CD45+ CD34- haematopoietic cells, megakaryocytes are identified as CD41+ CD42+ cells; macrophages as CD41- CD42- CD68/CD163+ cells; myelomonocytic cells as CD41- CD42- CD68/CD163- CD11b/CD14+ cells. The remaining negative population is further characterised to identify CD117/FceR+ cells as Eo/Baso/Mast cells. (c) Gating strategy for lymphoid panel. CD45+ lineage- haematopoietic cells are gated from live singlets. Then NK cells are identified as CD56+; CD7+ T progenitors as CD56- CD7+ CD5- and CD7+ CD5+ T progenitors as CD56- CD7+ CD5+. (d) Gating strategy for stromal panel. From live singlets, stromal cells are identified from non-erythroid (CD235a-) and non-haematopoietic (CD45-) cells. Endothelial cells are identified as CD31+, with a subset further defined as CD144+ vascular endothelial cells. MSCs/fibroblasts are identified as CD31- CD90+ CD271-, while CD31- CD90+ CD271+ cells as classified as MSCs. From the CD271+ MSCs, osteolineage cells are identified as ALP+ and LEPR+ MSCs are identified as LEPR+.

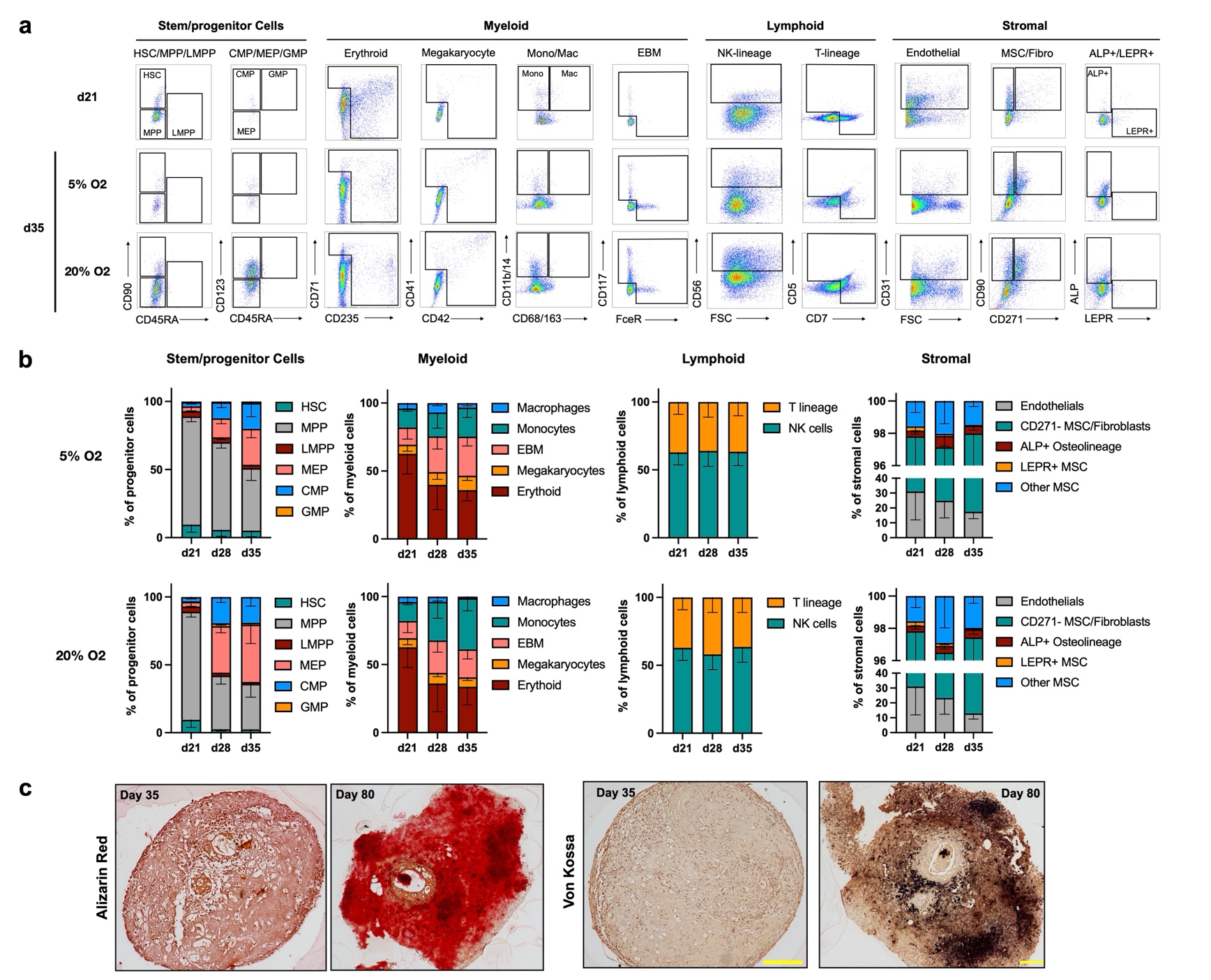

**Supplementary Figure 3: (a)** Representative flow cytometry plots at dat 21 and day 35 of comBO cultures maintained at either 5% or 20% oxygen from day 21-day 35. **(b)** Quantification of cell fractions at day 35 in either 5% or 20% oxygen between days 21 and day 35. No significant differences in populations are identified across conditions. **(c)** Assessment of mineralization of organoids at day 35 and 80 of cultures. Both alizarin red and von kossa staining confirm extensive mineralisation at day 80 of culture. scale bar = 200 µm.

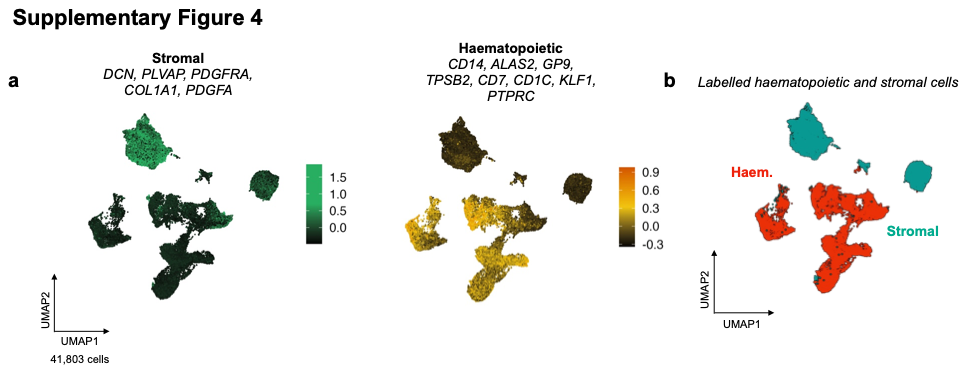

**Supplementary Figure 4: (a)** Total integrated day 20 single cell RNA sequencing data was first annotated into stromal and haematopoietic cells on the basis of canonical gene scoring (*DCN, PLVAP, PDGFRA, COL1A1, PDGFA)* for stromal cells and *CD14, ALAS2, GP9, TPSB2, CD7, CD1C, KLF1, PTPRC* for haematopoietic cells). **(b)** Objects were then subsetted on the basis of either stromal or haematopoietic lineages before further sub clustering and analysis.

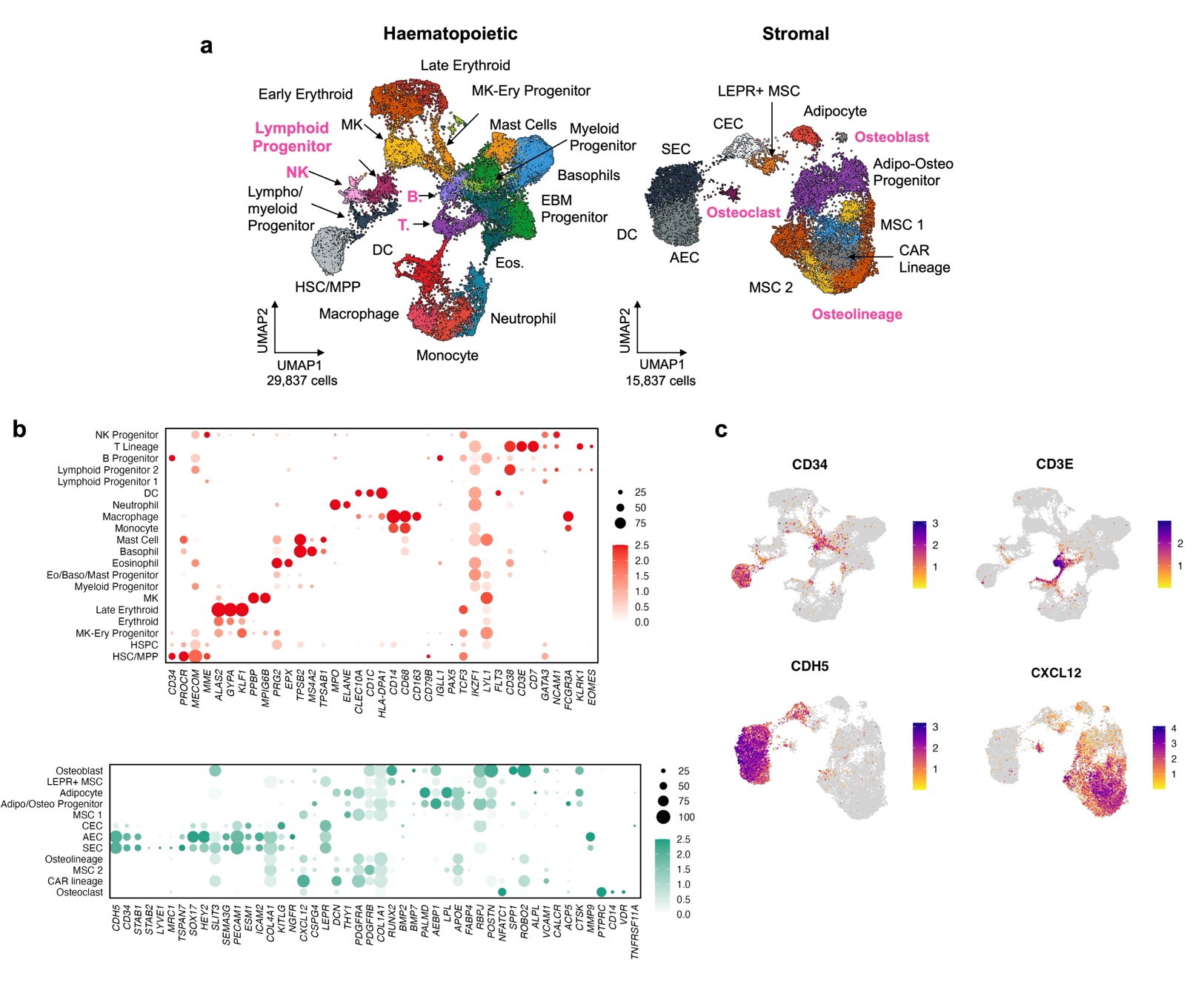

**Supplementary Figure 5:** Annotation of comBO UMAP using canonical genes. **(a)** An integrated haematopoietic object single RNA sequencing data set comprised of day 20 and day 35 comBO cells. Cells were collated from 4 independent differentiations which were cryopreserved prior to thawing and live cell sorting, and subsequent 10X preparation. Post quality control and pre-processing to separate haematopoietic and stromal cells, samples were annotated on the basis of canonical gene expression. **(b)** Haematopoietic cells included HSC/MPP (*CD34, PROCR, MECOM, MME*), HSPC/Lympho-Myeloid progenitors (low *CD34, PROCR, MECOM, CD79B, PAX5, GATA3, PRG2, TPSB2*), Erythroid lineage (*ALAS2, GYPA, KLF1, GFI1B)*, MK lineage (*GP9, PF4, PPBP, MPIG6B*), Eo/Baso/Mast (*GATA2*, *HDC, PRG2, TPSB2*), DC (*CD1C*, *HLA-DPA1*), monocyte/macrophage (*CD14, CD68, CD163*), B-progenitor (*CD79B, IGLL1*), neutrophil (*MPO, ELANE*) and T-lineage (*CD3E, CD7*) clusters were identified. Stromal cells included: Pan-endothelial genes *CDH5* and *PECAM1*, sinusoidal endothelial cells were identified by a cluster specific expression of *STAB2, LYVE1, TSPAN7* and *GGT5*. Arterial endothelial cells were separated by the lack of sinusoidal markers, and a high expression of *SOX17*. Capillary like endothelium were identified by high *KITLG*, *ESM1* expression and a lack of sinusoidal endothelial markers. MSCs were labelled with a canonical expression of MSC markers (e.g. *NGFR, CXCL12*), fibroblasts were separated by high *DCN* expression. Adipocytes were separated by a high expression of *AEBP1, PALMD, LPL, and APOE*. Osteoblasts were identified by a high expression of *RUNX2, POSTN, NFATC1, SPP1, ROBO2, MSX2 and VCAM1.* **(c)** Feature plots of key haematopoietic gene sets *CD3E* (T progenitor) and*CD34* (HSC/MPP). Feature plots of key stromal gene sets *CDH5* (Endothelial), *CXCL12* (CAR and other MSC) compartment is comprised of much more extensive range of stromal cells, a heterogenous vasculature, and a comBO specific osteoblast population. **(i,j)** this is confirmed by differential abundance (MiloR) analysis.

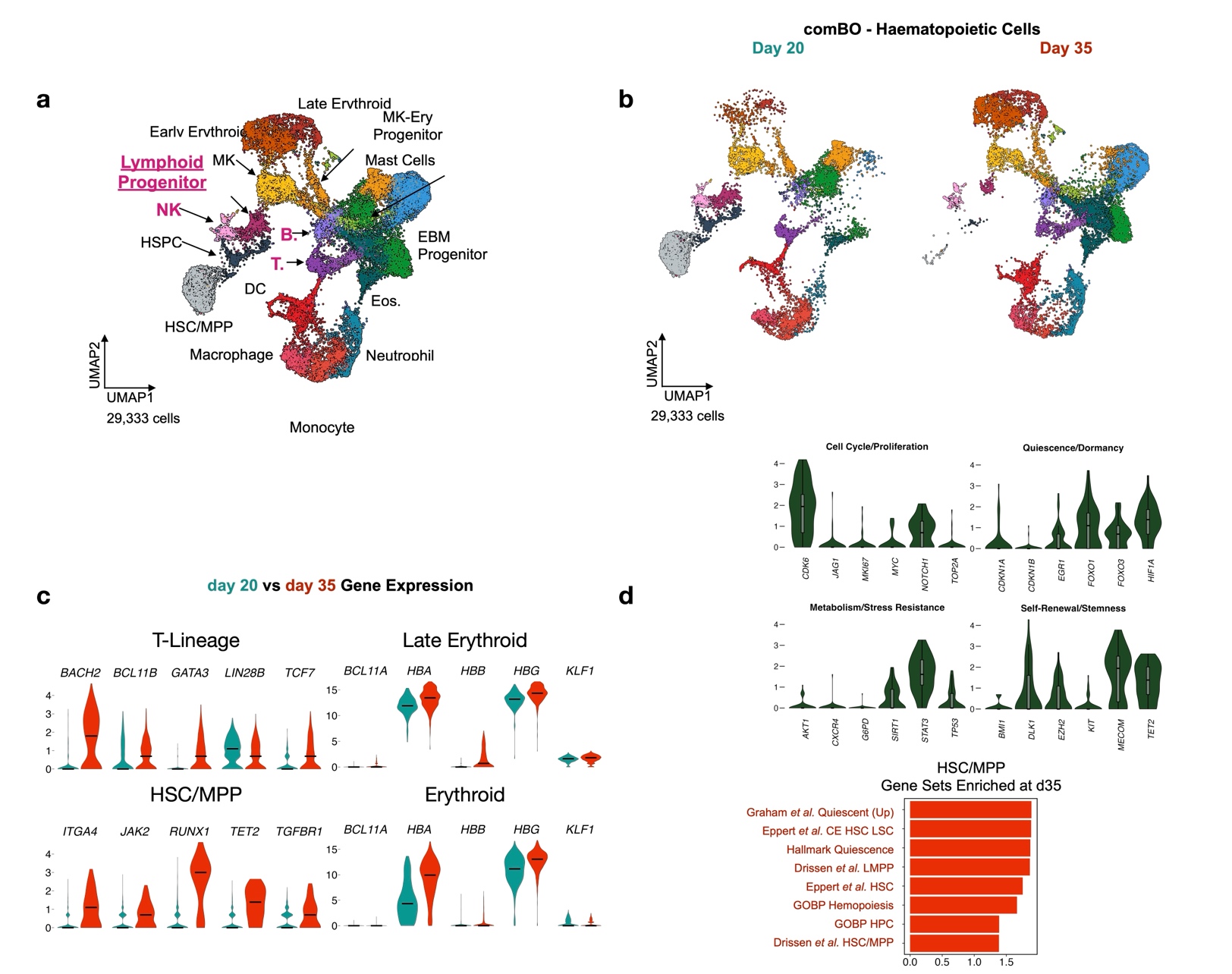
**Supplementary Figure 6: Integrated comBO d20 and d35 data.** Single cell RNA sequencing data of haematopoietic cell obtained at day 35 of comBO differentiation was integrated with day 20 data (reported in Figure 1) to investigate the maturation and development of haematopoietic cells over time. (a) A mix of lymphoid and myeloid cells at different stages of differentiation are observed (b) and these mature by day 35 including a particular loss of progenitor cells and expansion of committed progenitors, consistent with flow cytometry data. (c) A comparison of adult gene expression in key populations in day 20 and day 35 comBO. A notable increase in T-lineage markers of adult transcriptomic profiles is observed (*BACH2, BCL11B, GATA3, TCF7*) as welll as a reduction in the fetal marker *LIN28B****.*** HSC/MPPs demonstrated an increase in *ITGA4, JAK2, RUNX1, TET2, TGFBR1*, and erythroid cells demonstrate an increase in adult haemoglobin (*HBB)* at the late erythroid stage. The mix of *HBA* and *HBG* expression indicates a mixed fetal and adult transcriptome. (d) Further analysis of the HSC/MPP compartment indicates a decreased expression of cell cycle/proliferation markers (e.g, *TOP2A, MKI67*), the expression of markers of quiescence (*e.g. FOXO1/3, HIF1A*), and markers of self-renewal/stemness (e.g. *MECOM, TET2, DLK1*). Gene set enrichment analysis comparing day 20 and day 30 data against gene sets characterising adult, fetal, and diseased HSC/MPPs reported a significant up regulation in adult HSC profiles (*e.g.* Eppert et al. Drissen *et al.*) and hallmark quiescence gene profiles. Together this data indicates a trend towards adult gene expression in the comBO system.

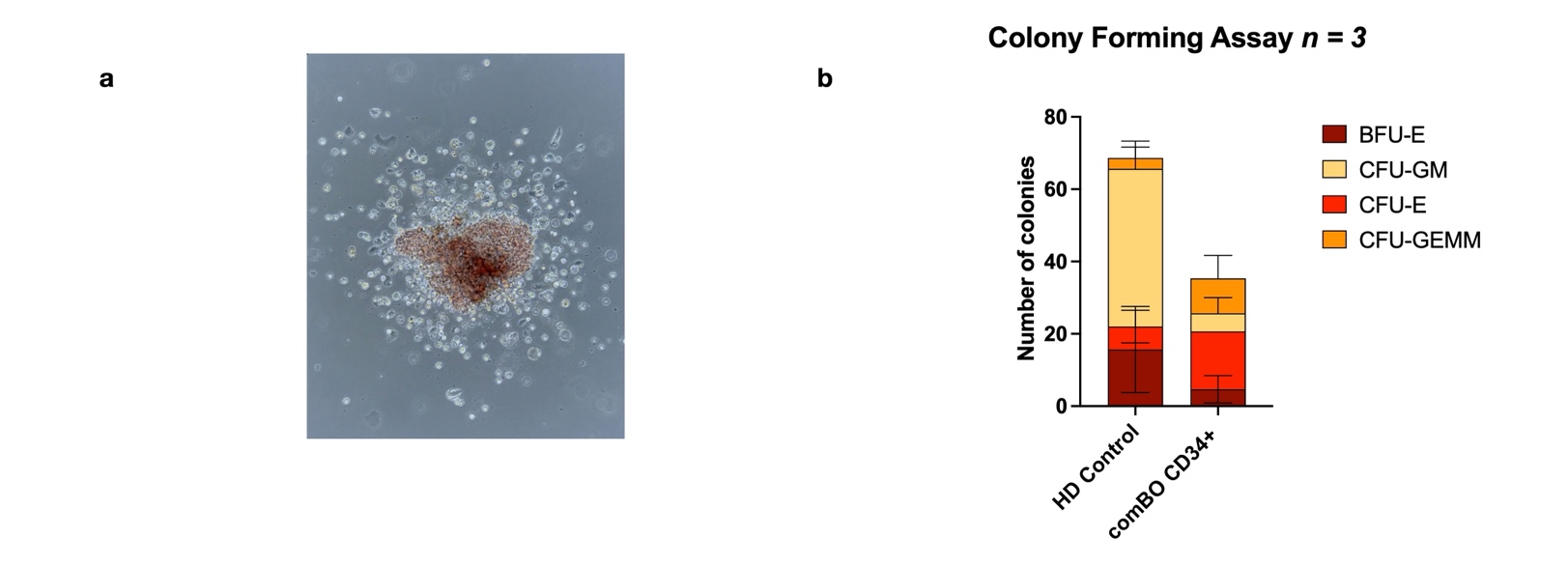

**Supplementary Figure 7:** A methocult assay was performed to assess the stem cell potential of CD34+ cells produced from the comBO. Cells were sorted on day 21 of differentiation (Live + CD34+ CD45+) and seeded in MethoCult at a density of 1,500 cells per well. In parallel, healthy donor CD34+ cells were similarly sorted and plated at an identical density. In total, 3 independent differentiations and healthy donors were used for this assay, with data summarised here. (a) Representative image of a colony forming unit. (b) Quantification of the number of colonies detected at day 14 of the methocult assay. While colonies were produced from comBO cells, they were notably fewer in number when compared to primary CD34+ healthy donors.

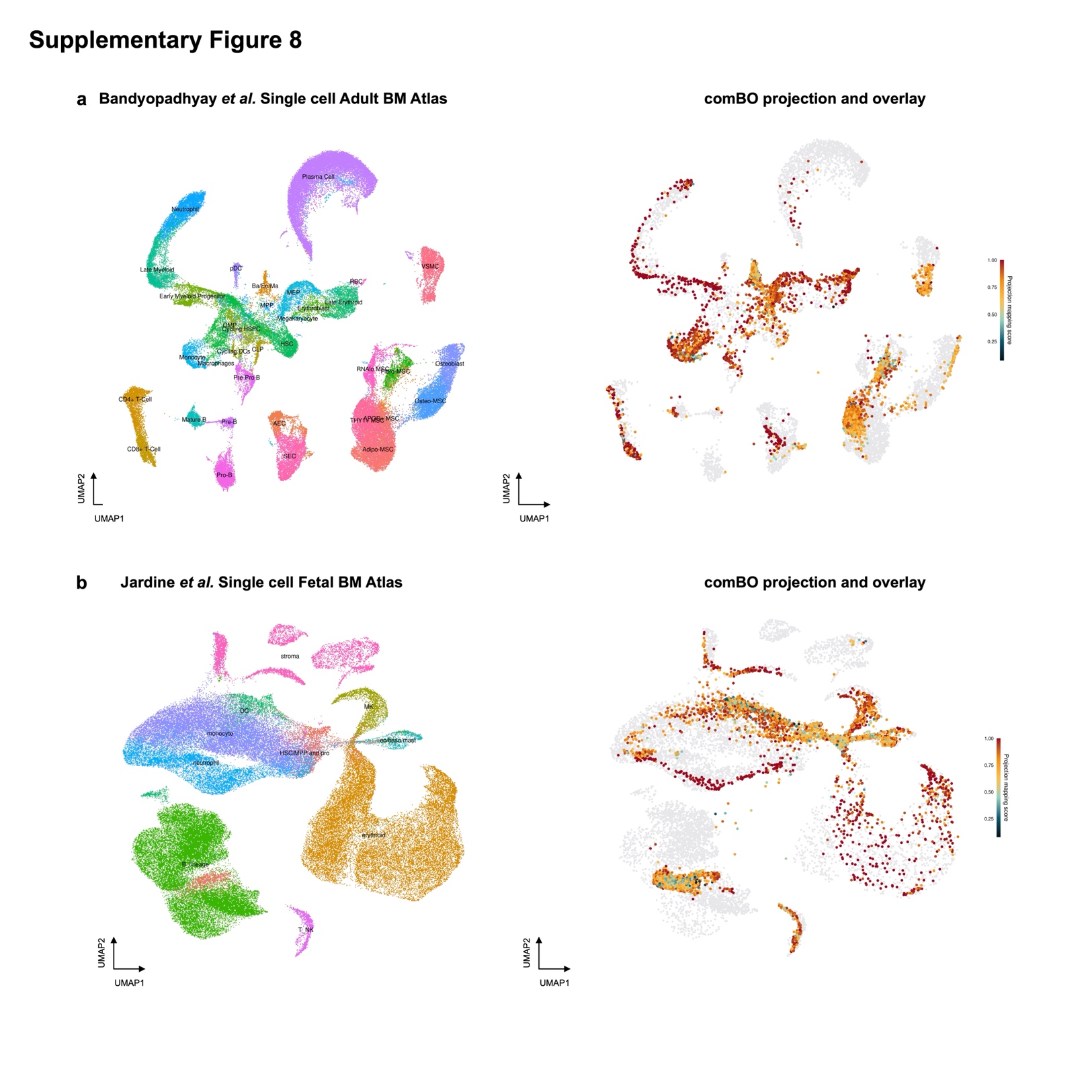

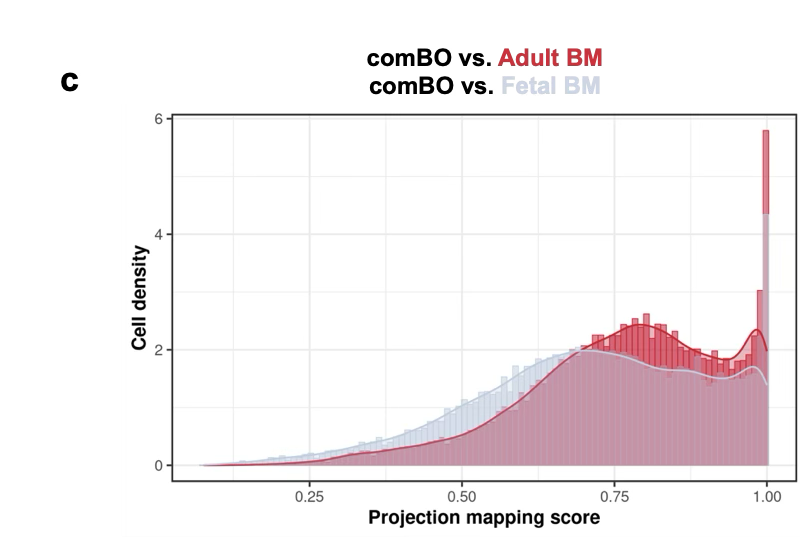

**Supplementary Figure 8:** Comparison of comBO (d35 data) and primary bone marrow single cell RNA sequencing data (adult and fetal, Bandyopadhay *et al.* and Jardine *et al.* respectively). (a) UMAP of a recently published adult bone marrow atlas, and (b) projection mapping of comBO scores after integration (analysis detailed in Materials and Methods). (c) UMAP of a primary fetal bone marrow data set capturing haematopoietic and stromal cells. (d) Projection scoring and overlay of similar cell populations. (d) A comparison of projection scores comparing comBO vs. Adult and fetal bone marrow demonstrating a higher degree of similarity (higher projection scores) in comparisons to adult bone marrow. Together, this data suggests a trend towards adult like gene expression.

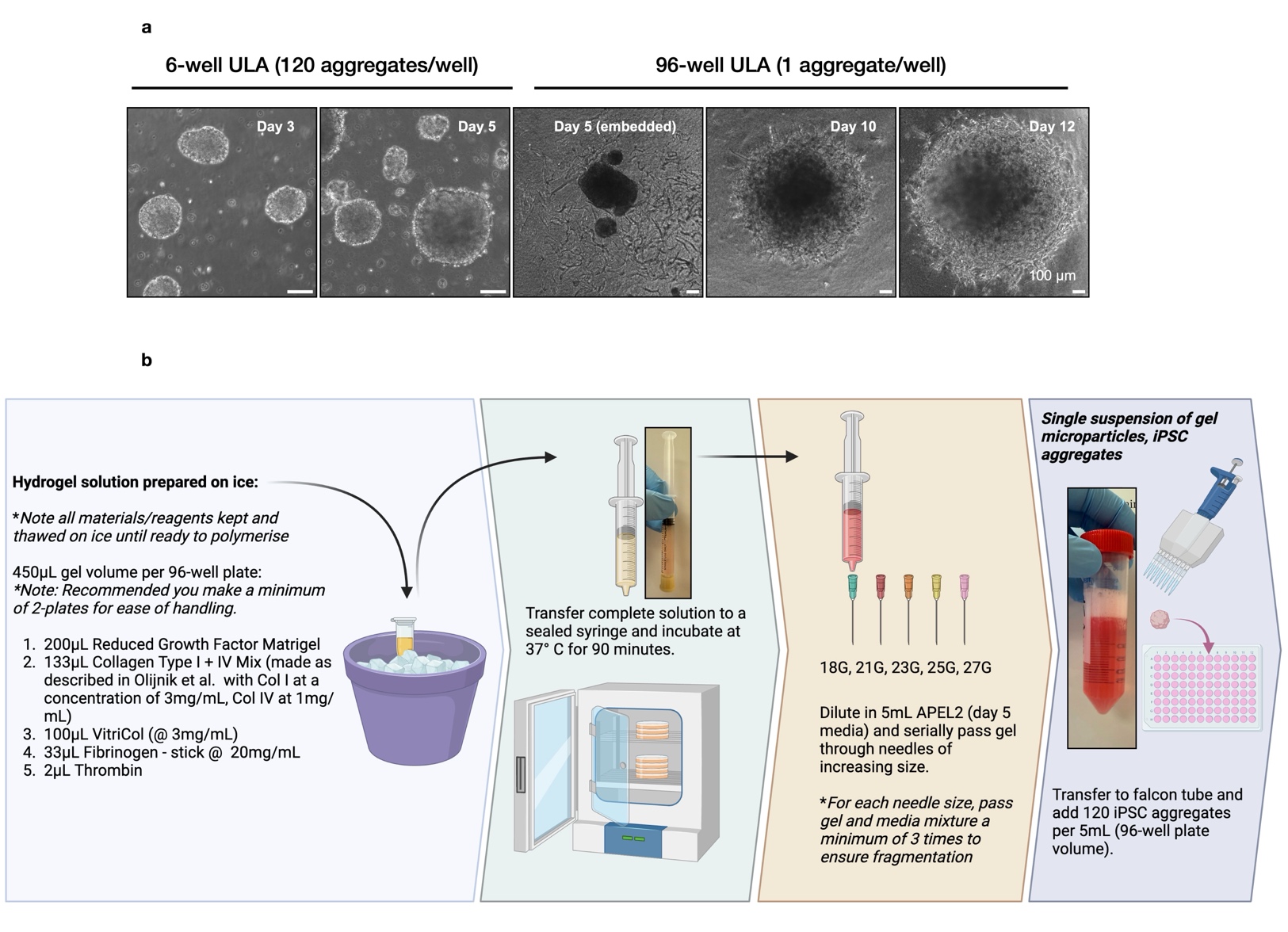

**Supplementary Figure 9:** Details of comBO generation using a granular microgel method. (a) Images of hiPSC aggregates in a 6-well ULA at days 3 and day 5, before embedding in a micro gel via (b) a step wise process. First a hydrogel solution is prepared on ice (as detailed in the diagram), before this solution is transferred to a sterile, sealed syringe. After incubation for 90 minutes, the fully polymerised hydrogel is diluted in 5mL of media (APEL2 supplemented with cytokines, knock out serum) and then passed serially through needles of increasing gauges to fragment the bulk gel and produce a suspension. Finally, the granular micro gel is transferred to a falcon tube, day 5 hiPSC aggregates are added to said tube, and then redistributed to a 96-well plate (50 μL volume per well). Plates are centrifuged (300G, 3min, room temperature) to compact the micro gels, and organoids returned to routine cell culture incubation. Schematic created in Biorender.com.

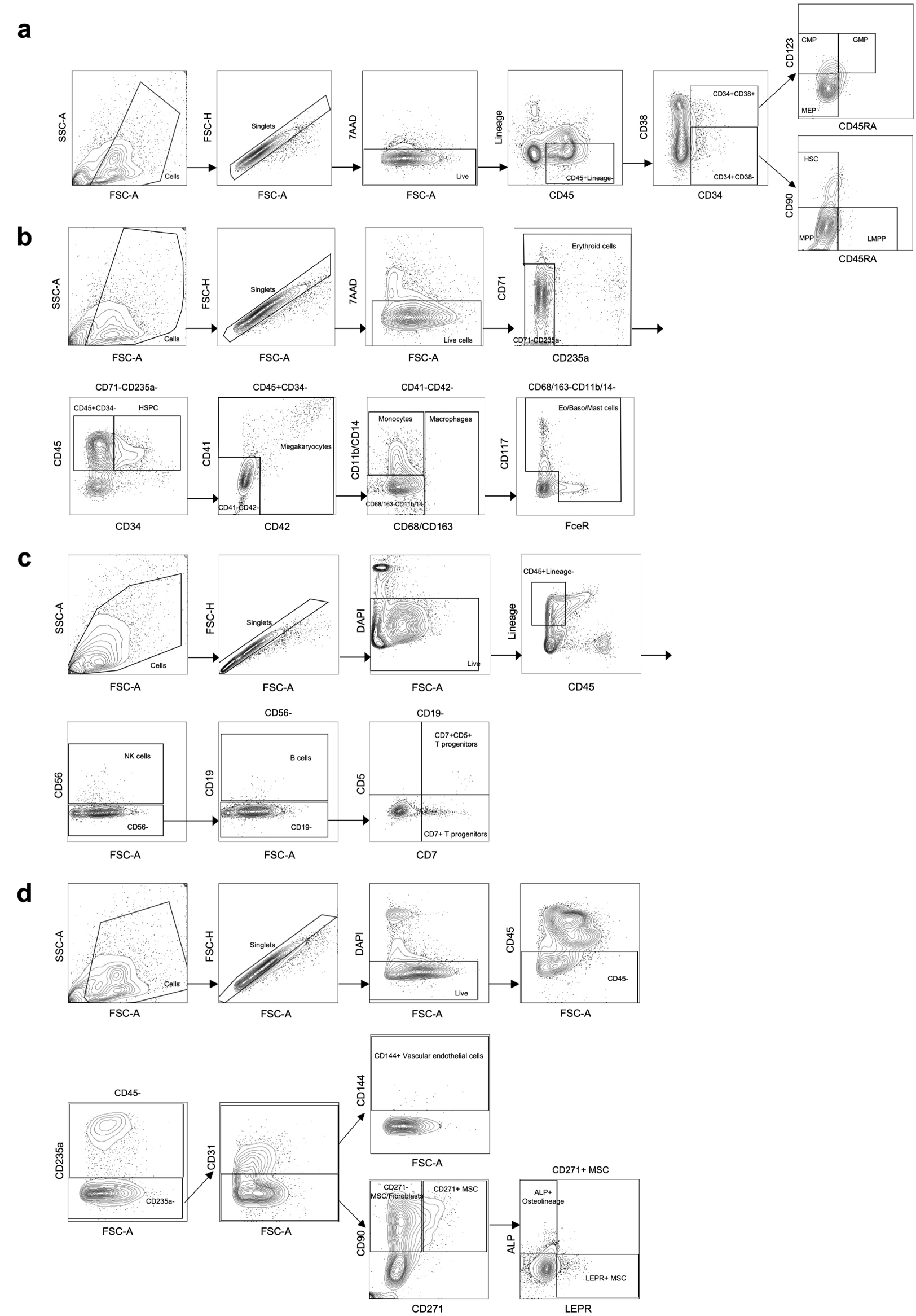

**Supplementary Figure 10:** Gating strategy for flow cytometry data reported in Fig. 2c-d. (a) Gating strategy for HSPC panel. Same as described in Supp. Fig 2a. (b) Gating strategy for myeloid panel. Same as described in Supp. Fig 2b. (c) Gating strategy for lymphoid panel. CD45+ lineage- haematopoietic cells are gated from live singlets. Then CD56+ cells are identified as NK cells, and CD56- cells are further analysed to identify B cells based on the expression of CD19. The CD19-cells are then used to identify T lineage cells. CD7+ CD5- cells as CD7+ T progenitors and CD7+ CD5+ cells as CD7+ CD5+ T progenitors. (d) Gating strategy for stromal panel. Same as described in Supp. Fig 2d.

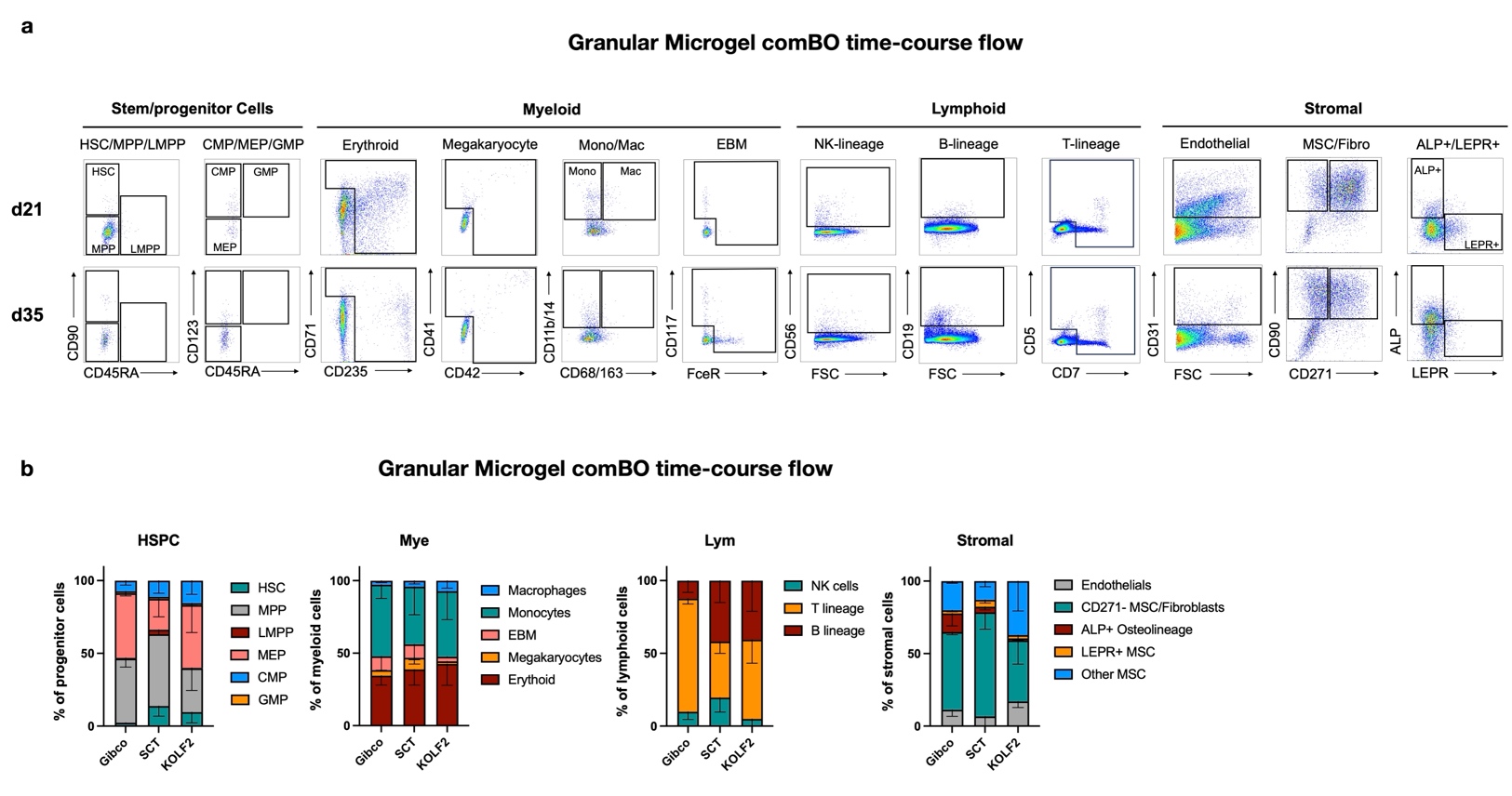

**Supplementary Figure 11:** Flow data generated by the granular microgel approach across time and multiple hiPSC lines. (a) Representative flow plots at day 21 and day 35 of comBO differentiation, demonstrating the differentiation of distinct lineages. B-lineage cells notably acquire an increase in CD19 between day 21 and day 35, while osteolineage cells for example, demonstrate an increase in ALP expression. (b) The comBO approach was applied across 3x distinct hiPSC lines showing a comparable generation of all lineages across each of the lines tested. Data is collated from 3x separate differentiations as 3x distinct biological repeats.

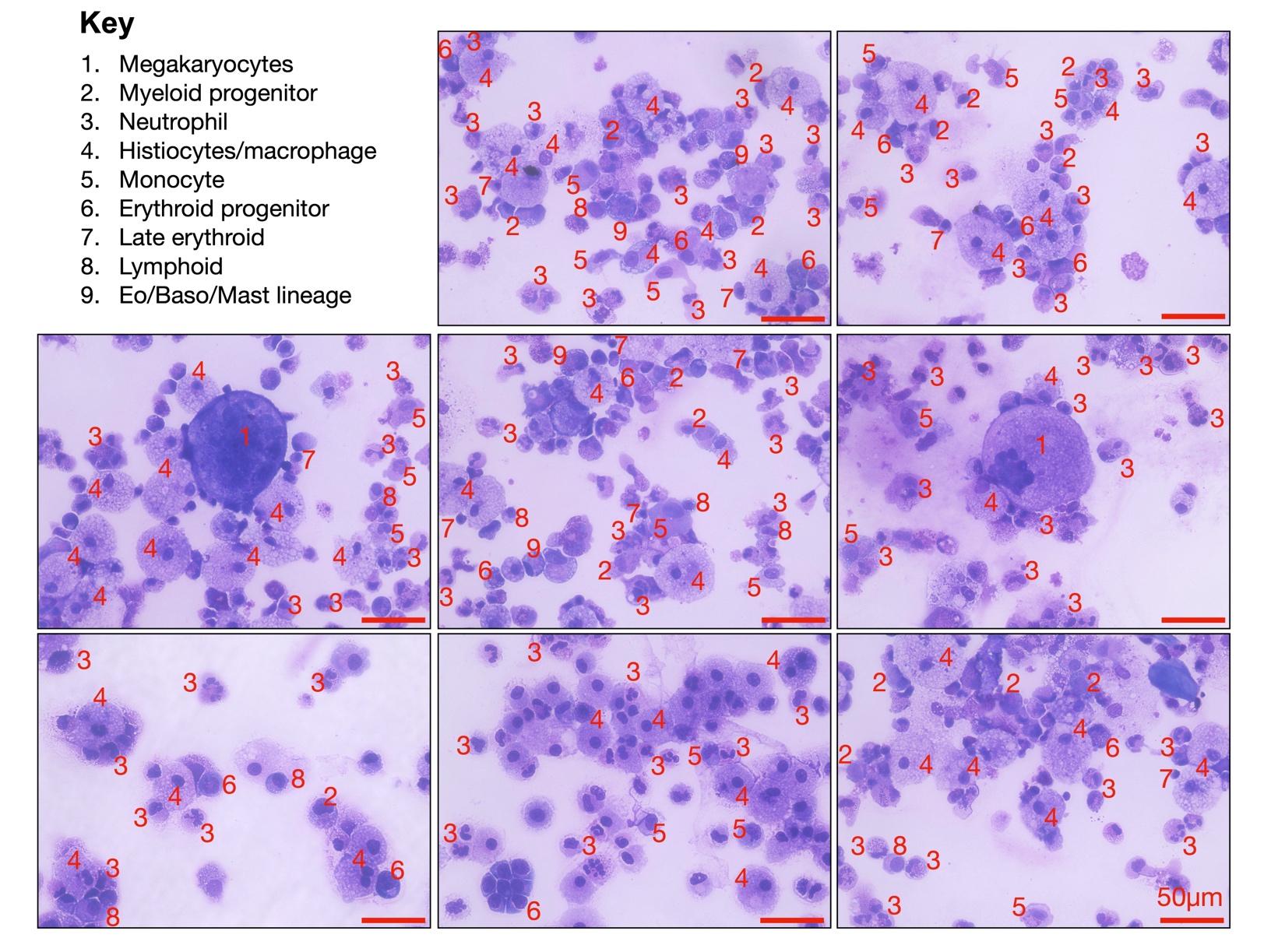

**Supplementary Figure 12:** Cytospins of comBO derived cells. comBOs were cultured under gentle agitation to encourage the egress of haematopoietic cells. Released haematopoietic cells were collected and prepared using cytospin and Wright's staining. Cells were then imaged and annotated as shown here. A range of cell types at different stages of maturation are observed, including myeloid and lymphoid cells as indicated.

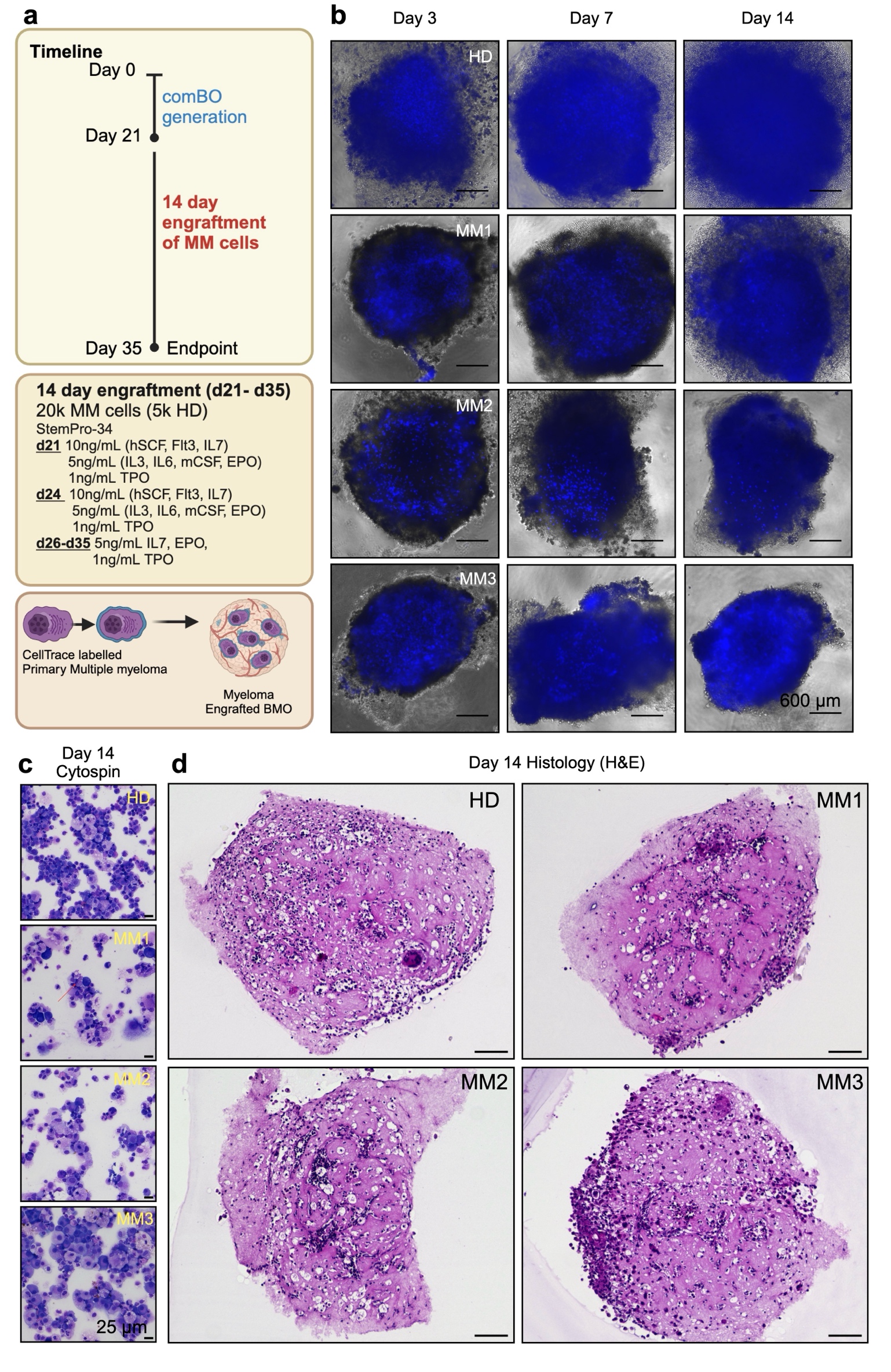

**Supplementary Figure 13:** Collated images from all 3 engrafted primary multiple myeloma comBO (MM-comBO) samples. (a) A workflow was devised whereby comBOs were generated using the microgel approach described and engrafted with CD138+ multiple myeloma cells (patients detailed in Supplementary Table 2). Patient cells were engrafted at 20,000 cells per well at day 21 and cultured for a further 2-weeks under cytokine conditions as detailed. (b) Myeloma cells were engrafted in parallel to healthy donor CD34+ cells and CellTrace labelled to track engraftment and proliferation over time. As expected, between days 3 and 14, healthy donor cells demonstrate a marked dilution of CellTrace consistent with proliferation and differentiation. Myeloma cells across all patients (MM1-MM3) demonstrate effective engraftment, but varying degrees of CellTrace dilution. (c) At day 14-post engrafted ganoids were dissociated and cytospin revealing viable, healthy myeloma cells post engraftment. (d) Organoids were also fixed and sectioned for H&E staining to confirm internalisation of myeloma cells across all patients used. Schematic generated by Biorender.com.

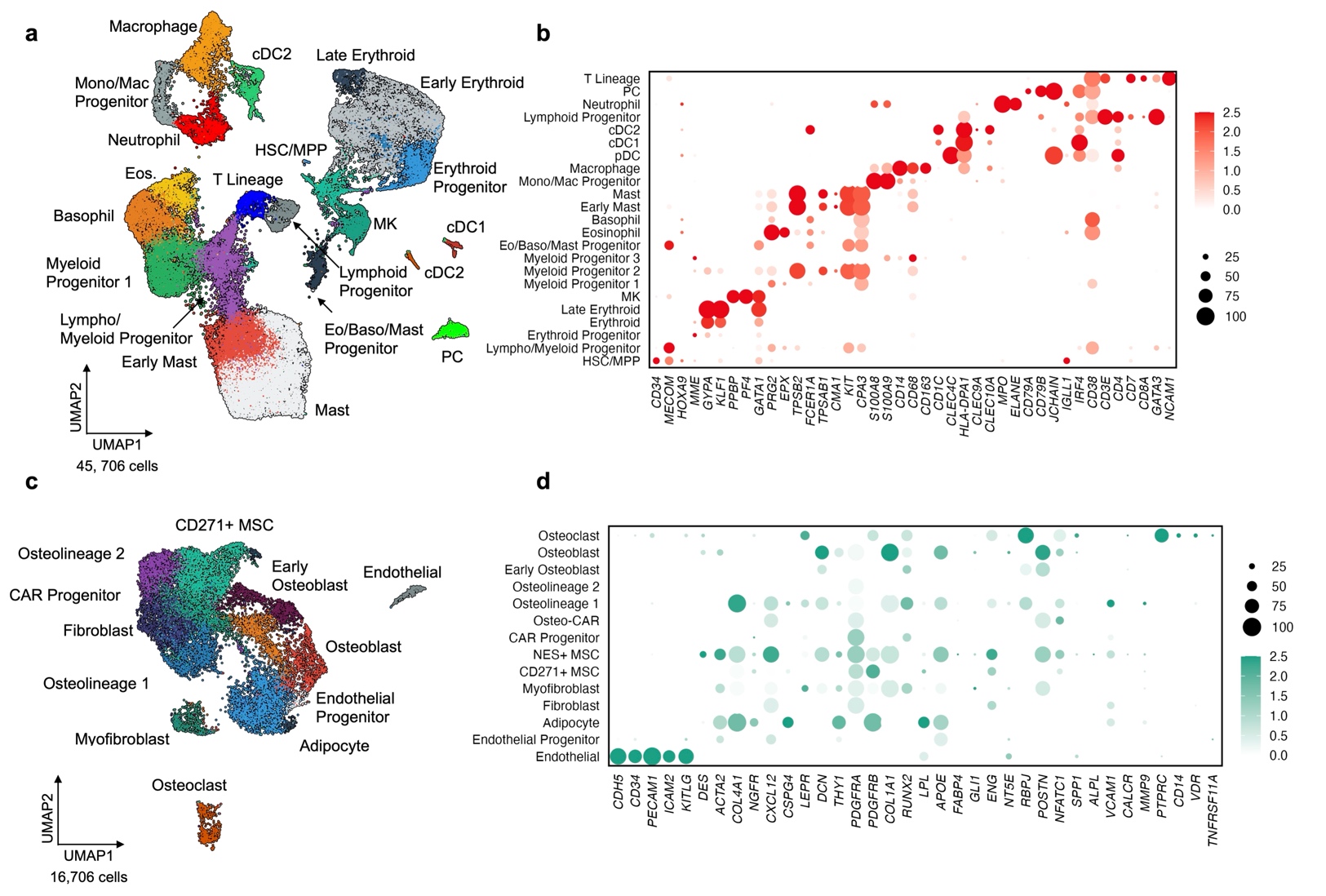

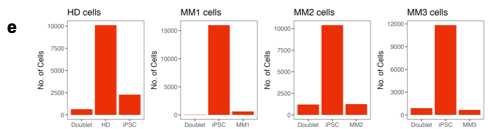

**Supplementary Figure 14:** Summary single cell data of unengrafted, control CD34+ healthy donor cells, and multiple myeloma engrafted comBOs (*n =* 3). (a) In total 45,706 cells were annotated as haematopoietic and defined by (b) canonical gene expression. (c) 16,706 stromal cells were collected post-processing, and annotated by (d) canonical gene expression. (e) Specific genotypes for engrafted samples were identified using souporcell.

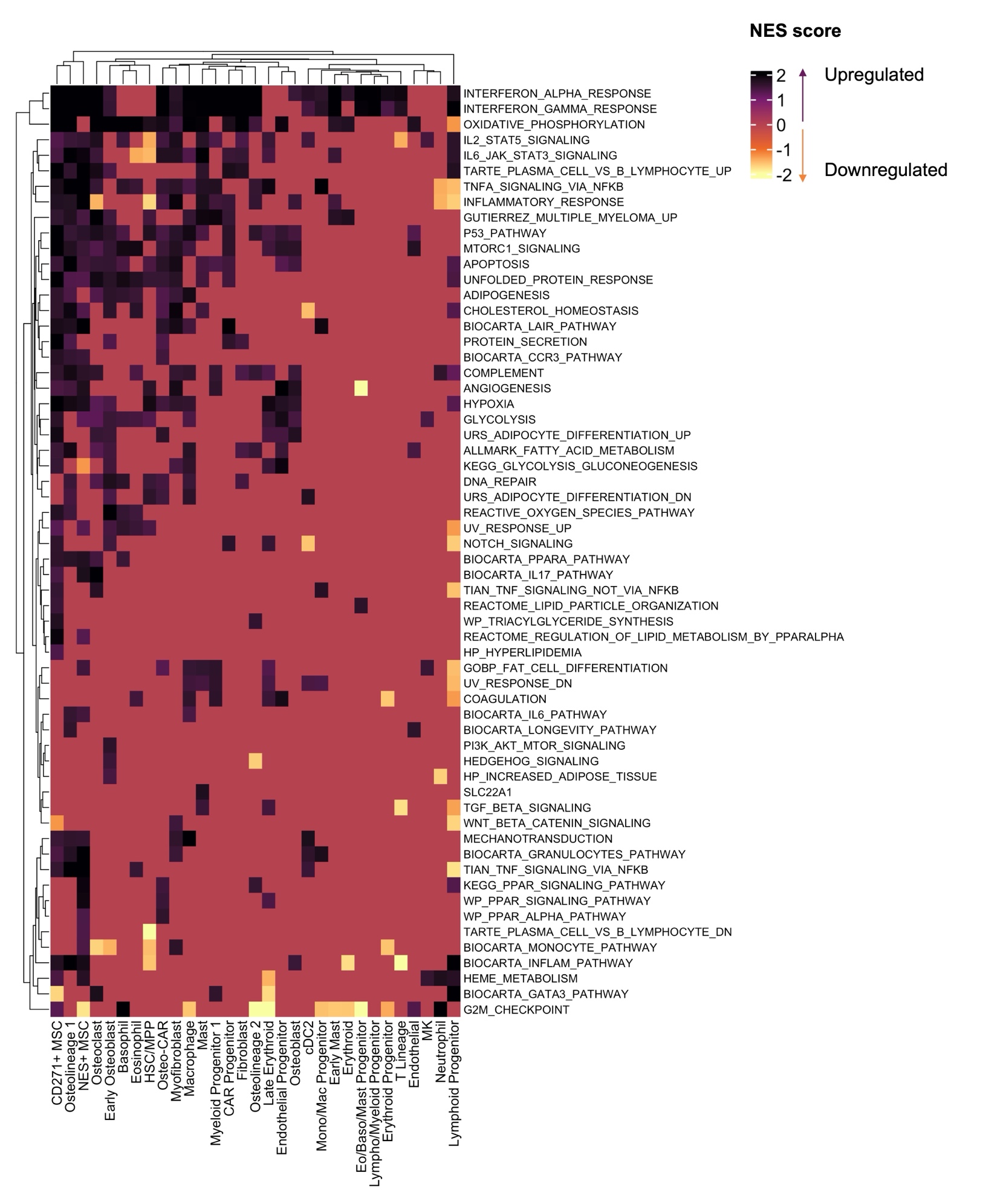
**Supplementary Figure 15:** A clustered heatmap of all significantly enriched gene sets in MM-comBO single cell RNA sequencing data (Normalised Enrichment Scores (NES)). Differential gene expression and gene set enrichment analysis were performed on an integrated control and multiple myeloma patient engrafted samples. CD271+ MSC, Osteolineage1, NES+MSC, osteoblasts, and osteoclasts cluster distinctly demonstrating dysregulation in interferon, IL2, IL6, TNF, and adipogenesis/fatty acid metabolism gene sets.

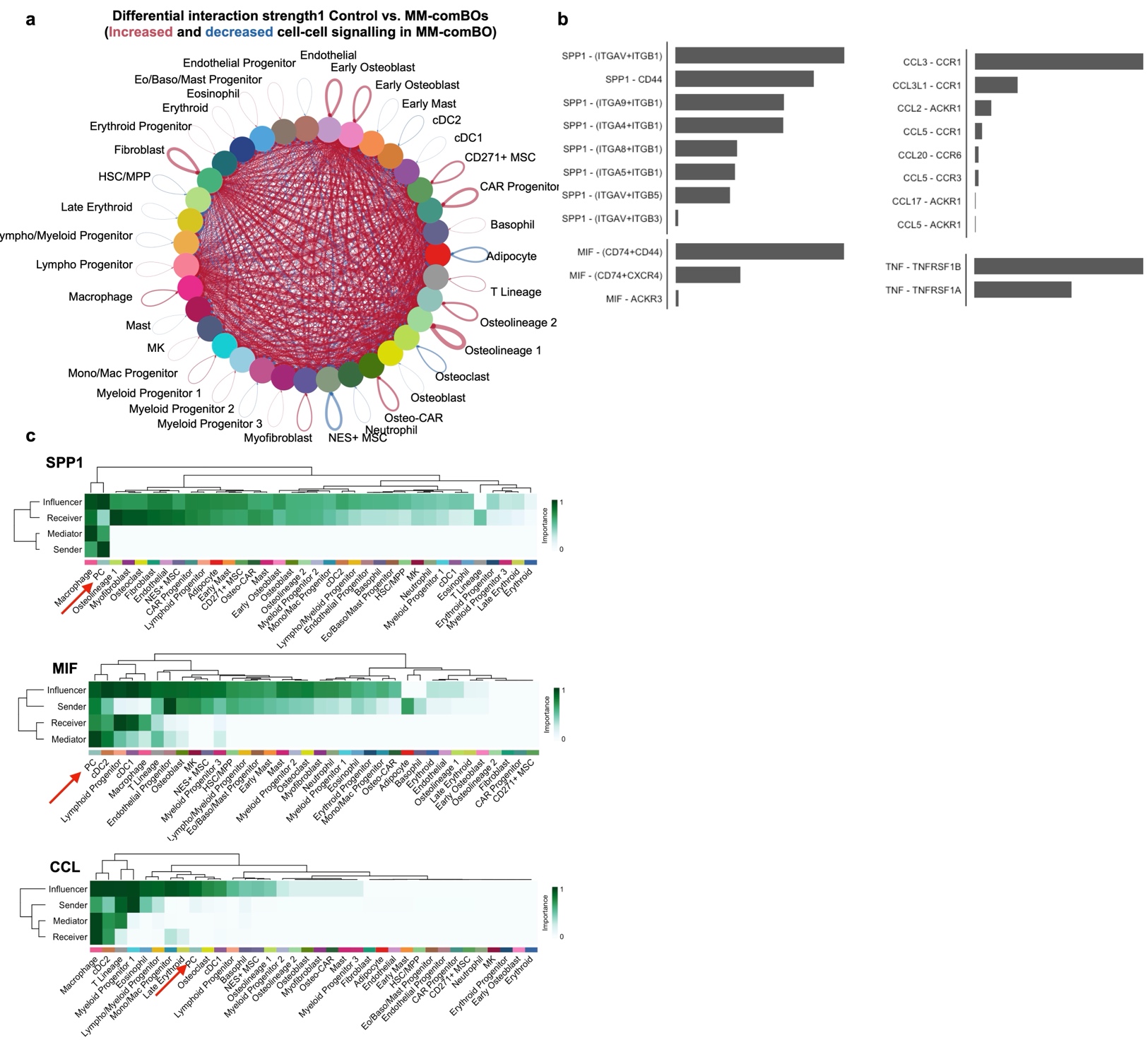

**Supplementary Figure 16:** Disrupted cell-cell interaction in MM-comBO. Cell-Cell interaction analysis (a) Circos plot of differential interaction strength, with red lines indicating predicted cluster-cluster interactions that are significantly upregulated in MM-comBO compared to controls. Blue lines indicate significantly downregulated interactions. (b) Summary of predicted ligand-receptor (LR) pairs in pathways identified in figure 6. SPP1-Integrin and SPP1-CD44, CCL3-CCR1, MIF-(CD74+CD44), and TNF-TNFRSF1B are the most significant interaction pairings in these pathways. (c) Heatmap of probable ‘senders’ of SPP1, MIF, CCL signals in MM-comBO identify PCs as the major source of SPP1 and MIF signalling that is dysregulated in MM-comBO.

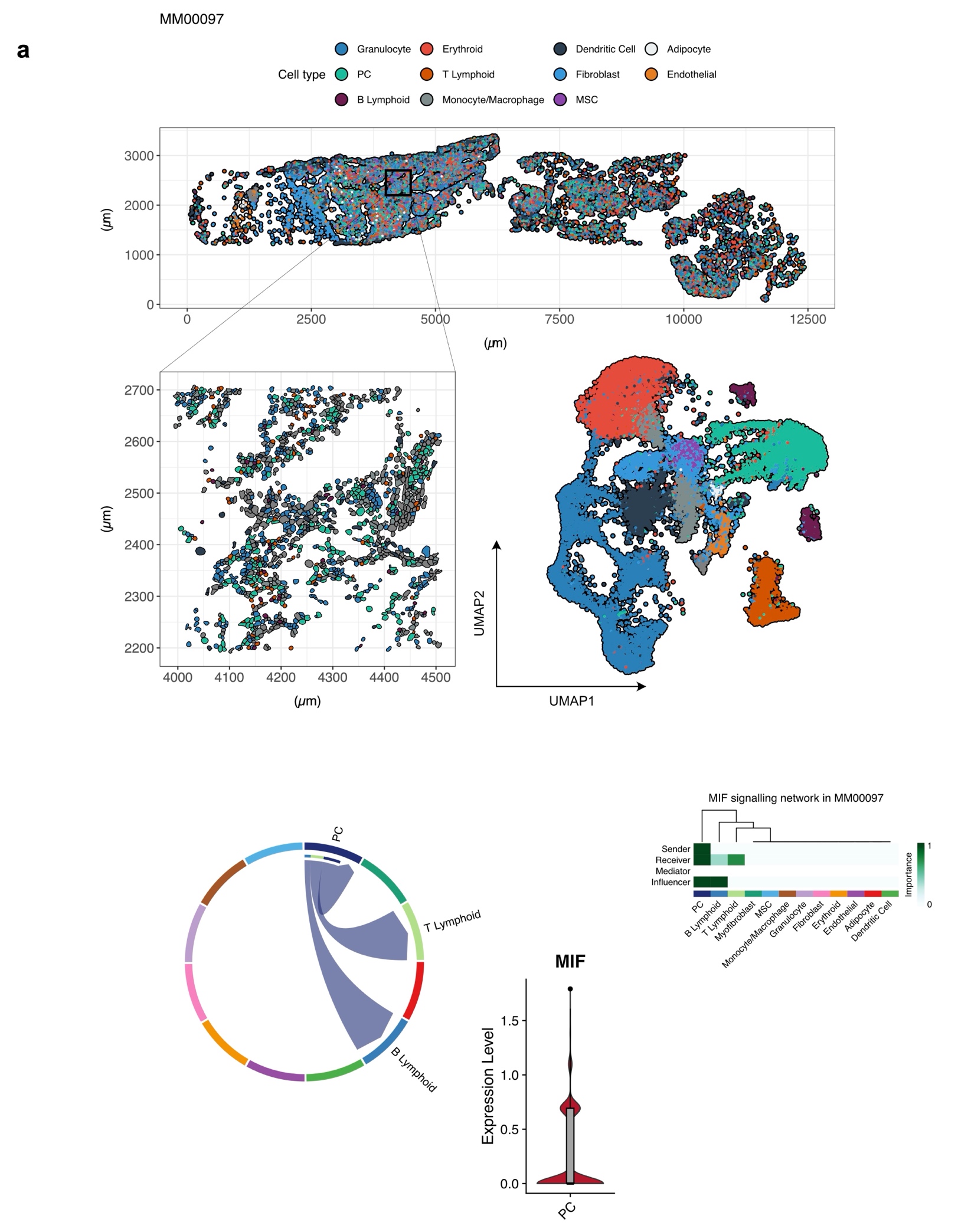

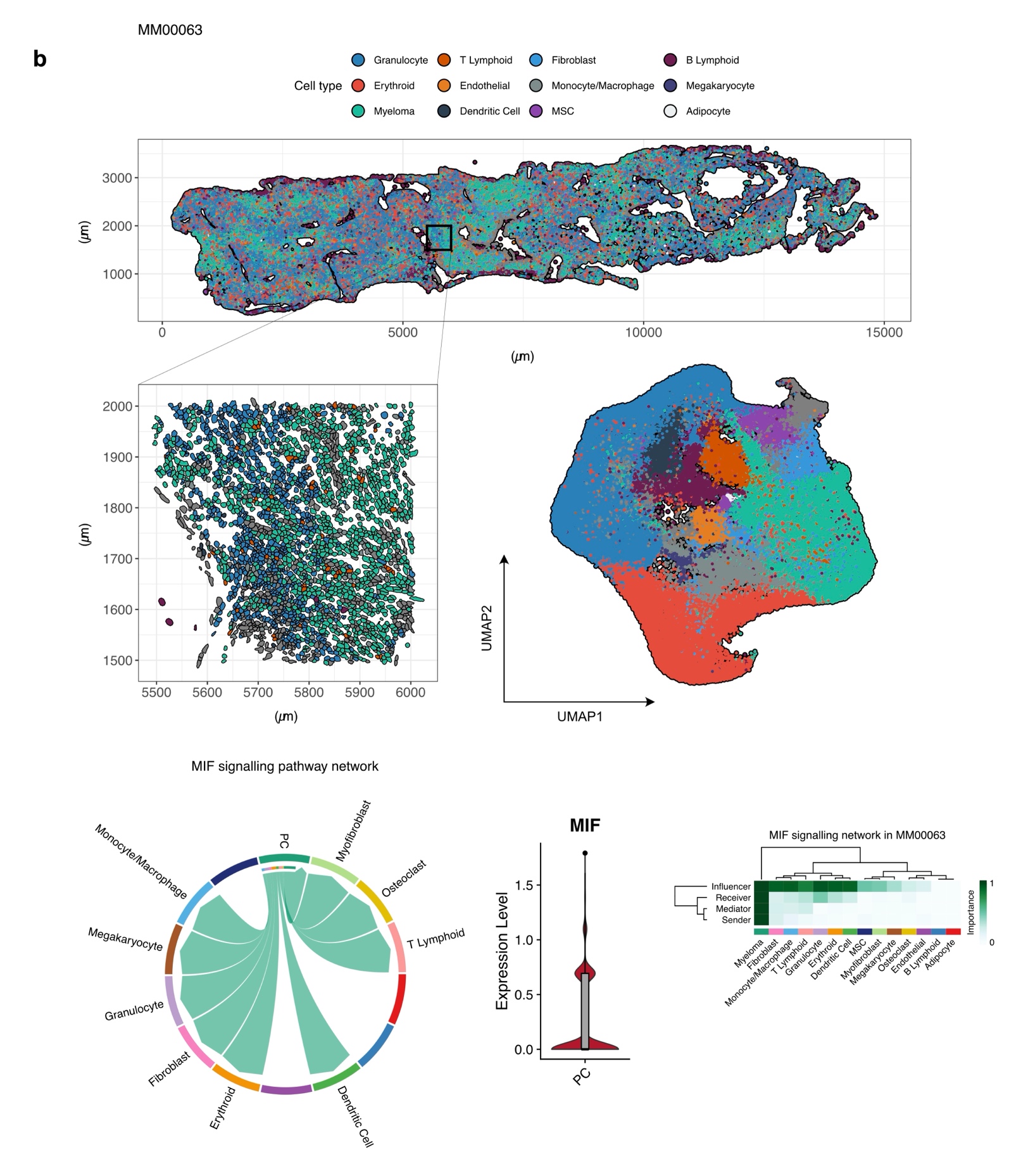

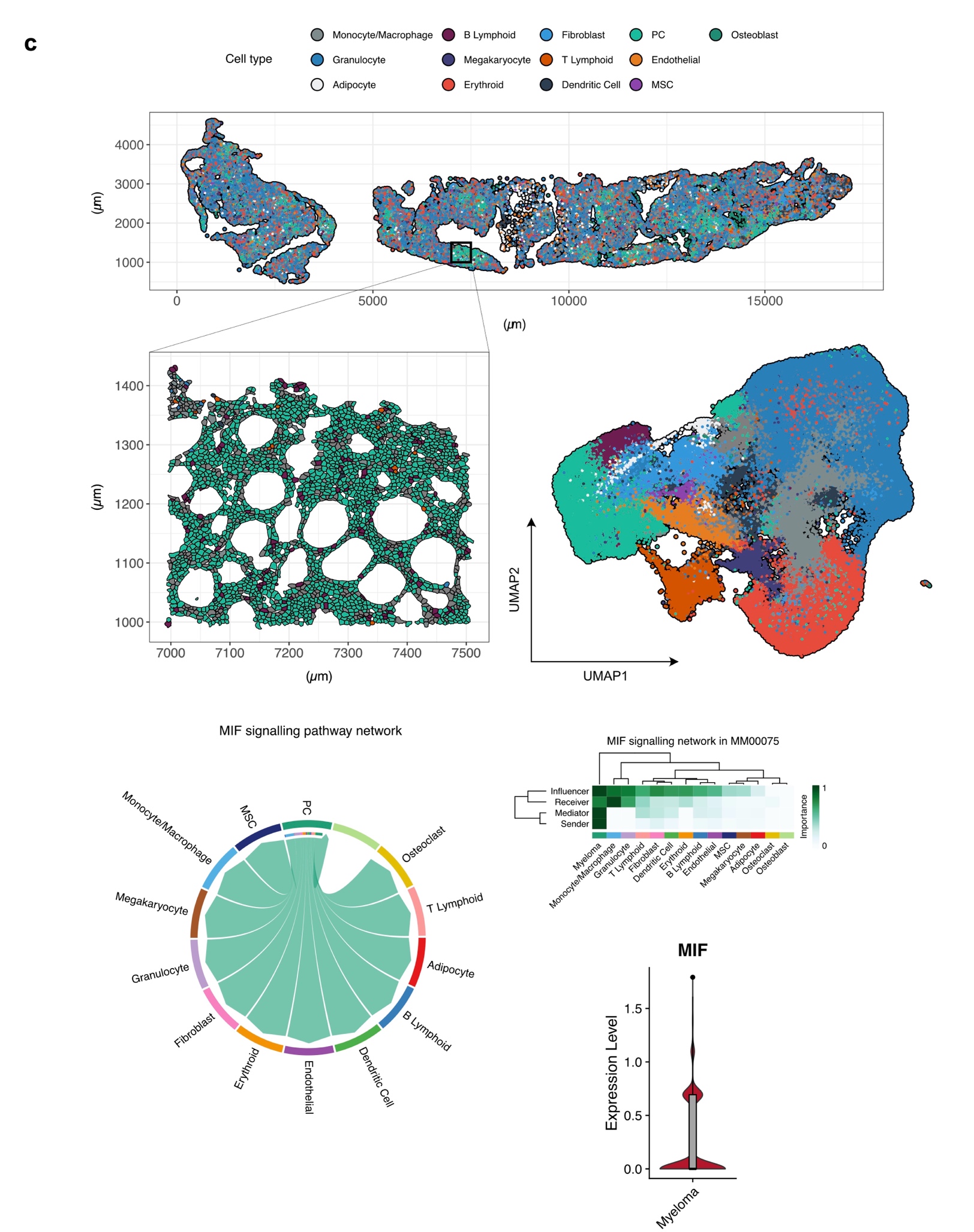

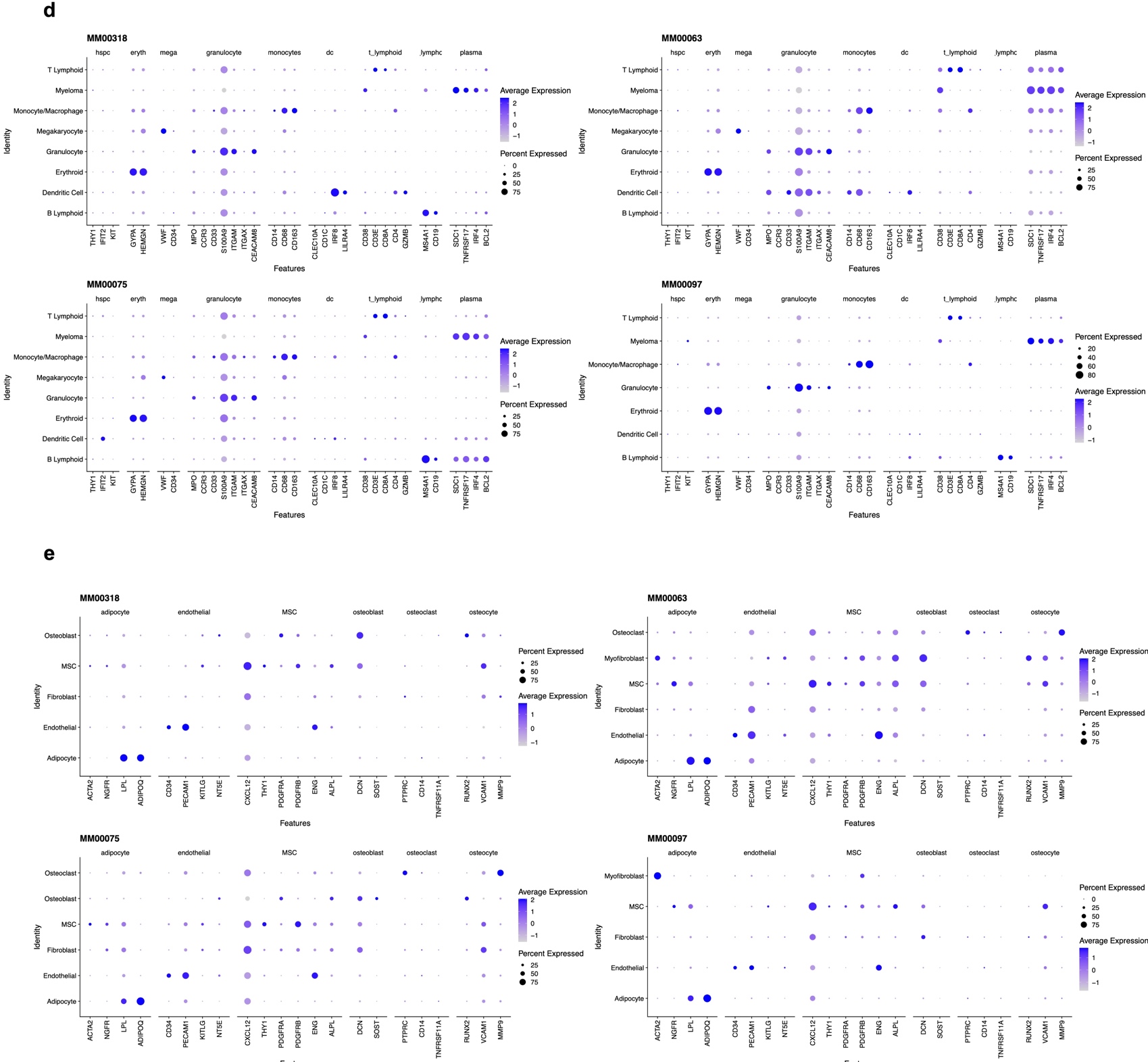

**Supplementary Figure 17:** Xenium spatial transcriptomic assessment of 3 additional multiple myeloma patients (Fig. 5d), patient details in Supplementary Table 2. (a-c) image, UMAP, and MIF signalling pathways shown per patient, with canonical gene expression (d) highlighting basis of annotation.

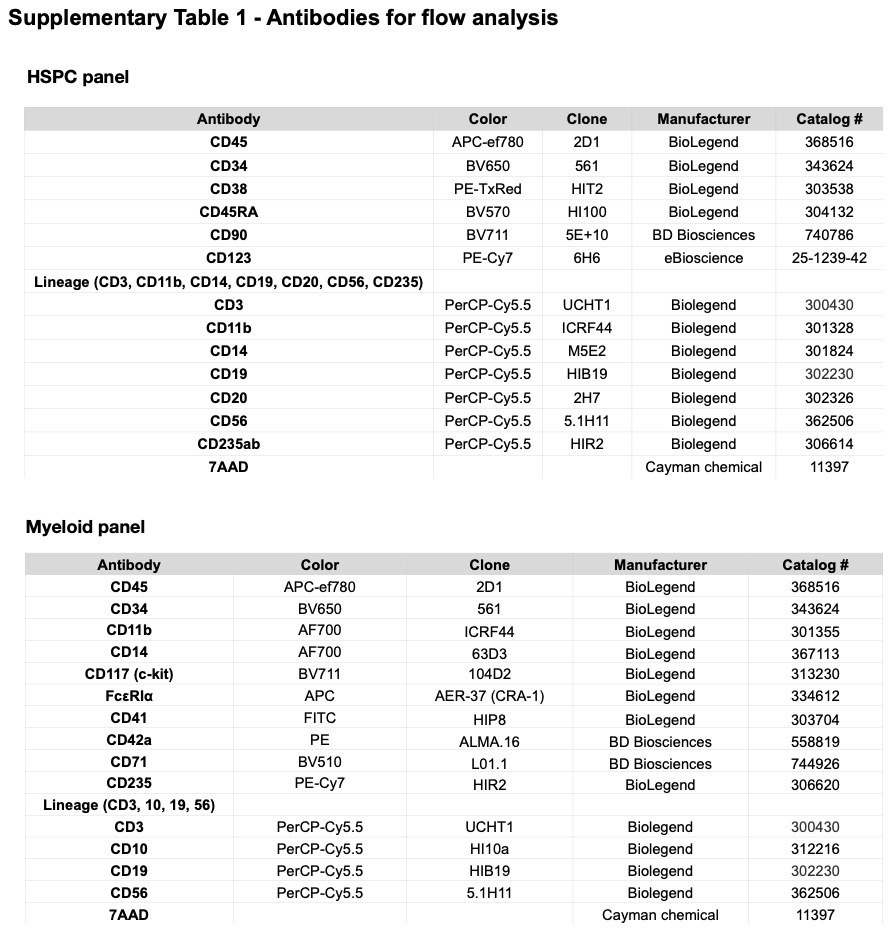

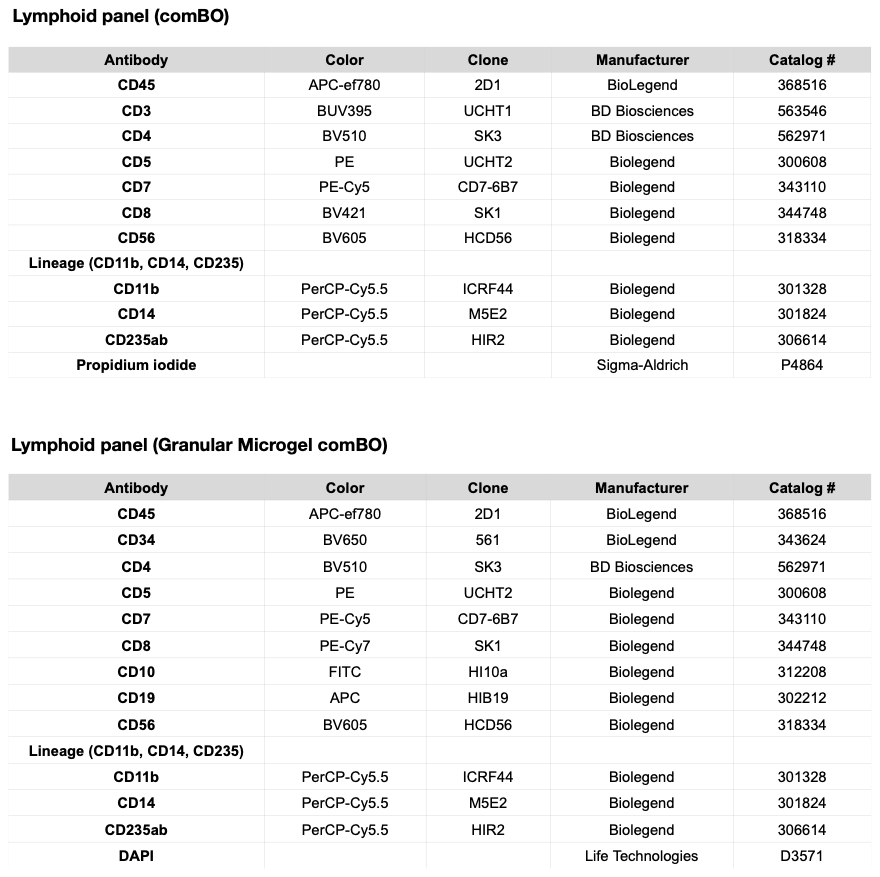

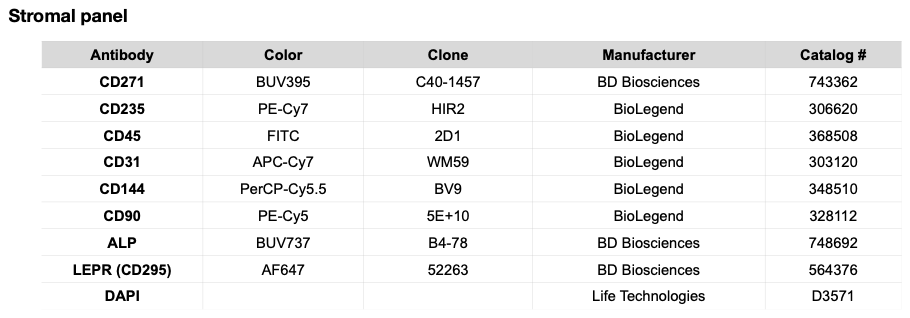

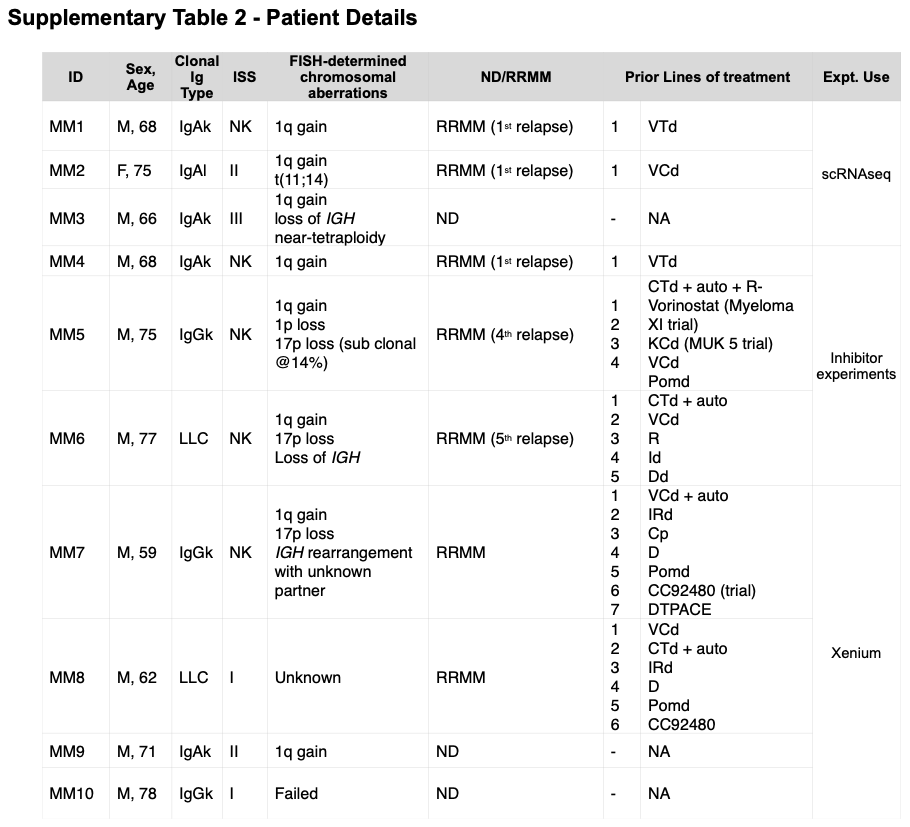

**Supplementary Table 2:** Clinical details of patients who donated cells or tissues used in experiments reported. Ig: immunoglobulin. ISS: International staging system for myeloma. FISH: Fluorescence in situ hybridisation. ND: newly diagnosed myeloma. RRMM: relapsed/refractory myeloma.

Myeloma clonal Ig types: IgA/G, k/l: myeloma expresses immunoglobulin A or G respectively, kappa or lambda light chain. LLC: lambda light chain myeloma.

NK: Not known. NA: Not applicable. *IGH:* immunoglobulin heavy chain gene.

Treatment regimens: VTd: velcade thalidomide dexamethasone; VCd: valcade cyclophosphamide dexamethasone; CTd: cyclophosphamide, thalidomide dexamethasone; auto: high dose melphalan autologous stem cell transplantation; R-vorinostat: lenalidomide, vorinostat; KCd: Carfilzomib, cyclophosphamide, dexamethasone; Pomd: pomalidomide, dexamethasone; R: lenalidomide; Id: Ixazomib, dexamethasone; Dd: Daratumumab, dexamethasone; IRd: Ixazomib, lenalidomide, dexamethasone; Cp: cyclophosphamide, prednisolone; D: daratumumab; DTPACE: dexamethasone, thalidomide, cisplatin, doxorubicin, cyclophosphamide, etoposide.

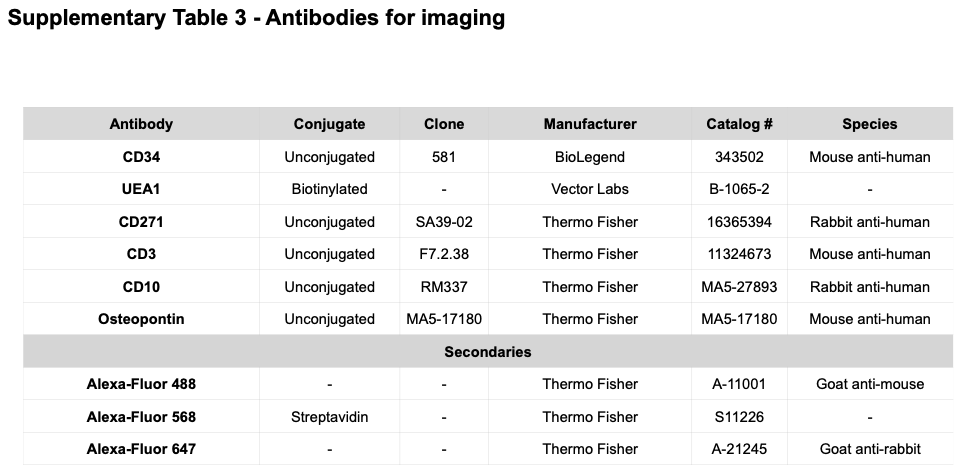
